# Primate heart regeneration *via* migration and fibroblast repulsion by human heart progenitors

**DOI:** 10.1101/2020.07.03.183798

**Authors:** Christine Schneider, Kylie S. Foo, Maria Teresa De Angelis, Gianluca Santamaria, Franziska Reiter, Tatjana Dorn, Andrea Bähr, Yat Long Tsoi, Petra Hoppmann, Ilaria My, Anna Meier, Victoria Jurisch, Nadja Hornaschewitz, Sascha Schwarz, Kun Lu, Roland Tomasi, Stefanie Sudhop, Elvira Parrotta, Marco Gaspari, Giovanni Cuda, Nikolai Klymiuk, Andreas Dendorfer, Markus Krane, Christian Kupatt, Daniel Sinnecker, Alessandra Moretti, Kenneth R. Chien, Karl-Ludwig Laugwitz

## Abstract

Human heart regeneration is one of the most critical unmet clinical needs at a global level^1^. Muscular regeneration is hampered both by the limited renewing capacity of adult cardiomyocytes^2-4^ and the onset of cardiac fibrosis^5,6^, resulting in reduced compliance of the tissue. Primate have proven to be ideal models for pluripotent stem cell strategies for heart regeneration, but unravelling specific approaches to drive cell migration to the site of injury and inhibition of subsequent fibrosis have been elusive. Herein, by combining human cardiac progenitor lineage tracing and single-cell transcriptomics in injured non-human primate heart bio-mimics, we uncover the coordinated muscular regeneration of the primate heart *via* directed migration of human ventricular progenitors to sites of injury, subsequent fibroblast repulsion targeting fibrosis, and ultimate functional replacement of damaged cardiac muscle by differentiation and electromechanical integration. Single-cell RNAseq captured distinct modes of action, uncovering chemoattraction mediated by CXCL12/CXCR4 signalling and fibroblast repulsion regulated by SLIT2/ROBO1 guidance in organizing cytoskeletal dynamics. Moreover, transplantation of human cardiac progenitors into hypo-immunogenic CAG-LEA29Y transgenic porcine hearts following injury proved their chemotactic response and their ability to generate a remuscularized scar without the risk of arrhythmogenesis *in vivo*. Our study demonstrates that inherent developmental programs within cardiac progenitors are sequentially activated in the context of disease, allowing the cells to sense and counteract injury. As such, they may represent an ideal bio-therapeutic for functional heart rejuvenation.

---

Whereas mammals do have the capacity to undergo endogenous cardiac regeneration during development and shortly after birth^7^, the regenerative capacity of the human heart in adulthood is markedly low^3^. The inability to replace lost myocardium is accompanied by extensive tissue remodelling and fibrosis, leaving patients with cardiac disease vulnerable to heart failure. Although numerous drugs and mechanical devices can moderately improve cardiac function, such approaches do not replace lost cardiomyocytes (CM) or abolish fibrotic scar formation. Over the past decade, *bio*-therapies have emerged as innovative strategies for heart repair^8-11^. Induction of endogenous CM proliferation^12-15^, *in vivo* direct reprogramming of non-CMs to a cardiac fate^16^, and exogenous transplantation of human pluripotent stem cell (hPSC)-derived CMs^17^ or cardiac progenitors^18^ have been recently explored as potential approaches to generate *de novo* myocardium.

Studies in lower vertebrates, where robust cardiac regeneration occurs throughout life, have demonstrated that endogenous heart repair is a highly coordinated process involving inter-lineage communication, cellular de- and re-differentiation, migration, and extracellular matrix (ECM) remodelling without fibrotic scarring^19-22^. Similar programs are the foundation of organ morphogenesis and are inherent of embryonic cardiac progenitors. During heart development, defined embryonic precursors, including first heart field (FHF) and second heart field (SHF), give rise to distinct cardiac compartments and cell types^23,24^. While FHF cells, marked by HCN4 and NKX2.5, are fated to differentiate early into CMs of the primitive heart tube, ISL1^+^ SHF has broader lineage potential and its differentiation is preceded by an extensive proliferation and directed migration into the forming myocardium^25-27^. We have recently reported the generation of an enriched pool of hiPSC-derived ISL1^+^ ventricular progenitors (HVPs), which can expand and differentiate into functional ventricular CMs *in vitro* and *in vivo*^28^. Here, we sought to determine whether HVPs could effectively provide primate heart regeneration by orchestrating and activating sequential programs of cardiac development, ultimately leading to *de novo* myocardium formation and fibrotic scar-less healing.

## HVPs functionally repopulate a chronic injury model of non-human primate heart slices

During cardiogenesis, heart progenitors develop in a three-dimensional (3D) micro-environment incorporating important cues from myocardial architecture and electromechanical forces. To dissect molecular steps of HVP-mediated cardiac repair at a single cell level, we established an *ex vivo* 3D chimeric model where HVPs were exposed to the complex structural and molecular environment of adult heart tissue from non-human primate (NHP). In customized bio-mimetic chambers^29^, native NHP left ventricle (LV) slices (∼300 μm thickness) were cultured under physiological preload (1 mN) and continuous stimulation (1 Hz pacing), allowing proper structural and functional preservation for up to 14 days (D) (Fig. 1a, b and Extended Data Fig. 1a). Subsequently, progressive tissue deterioration occurred, as indicated by the gradual loss of contractile force that coincided with increasing apoptotic CM death over time (Fig. 1b, c and Extended Data Fig. 1a, b), offering an ideal setting for investigating cell-based mechanisms of cardiac repair. *NKX2.5*^eGFP/*wt*^ human embryonic stem cells (hESC) were coaxed towards *ISL1*/*NKX2.5* expressing heart progenitors using our previously described two-step protocol that enriches for HVPs^28^, with small number of multipotent ISL1^+^ precursors^30^ (Fig. 1a and Extended Data Fig. 1c). After magnetic-activated cell sorting (MACS)-based depletion of undifferentiated hESCs, cells were seeded onto the NHP LV slices by standardized bioprinting (Extended Data Fig. 1c, d). eGFP expression regulated by the NKX2.5 locus allowed live tracing of HVPs and their derivative CMs within the NHP myocardium (Extended Data Fig. 1e, top). Labelling with EdU and activated caspase-3 (ClCasp3) indicated that eGFP^+^ cells were highly proliferative during the first 2 weeks, but stopped expanding by D21 when eGFP^-^ NHP-CMs underwent massive apoptosis (Fig. 1c and Extended Data Fig. 1f). This corresponded to the extensive differentiation towards CMs and downregulation of ISL1 expression (Extended Data Fig. 1g). Remarkably, heart slices gradually regained contractile force in the third week of co-culture, reaching 2mN force generation that was further maintained up to D50 (Fig. 1b and Extended Data Fig. 1e). Immunohistochemistry for atrial and ventricular muscle markers (MLC2a / MLC2v) revealed that the majority of eGFP^+^ cells acquired a ventricular muscle identity over time (Fig. 1d). By D21, ∼19% expressed both markers, indicative of CM entering the ventricular lineage, and ∼69% were already exclusively MLC2v positive, representing maturing ventricular CMs. The latter reached ∼81% by D50 (Fig. 1d). Moreover, at this stage, most eGFP^+^/MLC2v^+^ CMs presented a rod-shape appearance with well-aligned myofibrils, structural characteristics of mature muscle cells (Fig. 1d). A small proportion of human cells expressing the endothelial marker CD31 were also detected (∼14% on D21 and ∼7% on D50) (Fig. 1e), likely arising from multipotent precursors within the HVP pool.

**Figure 1.**
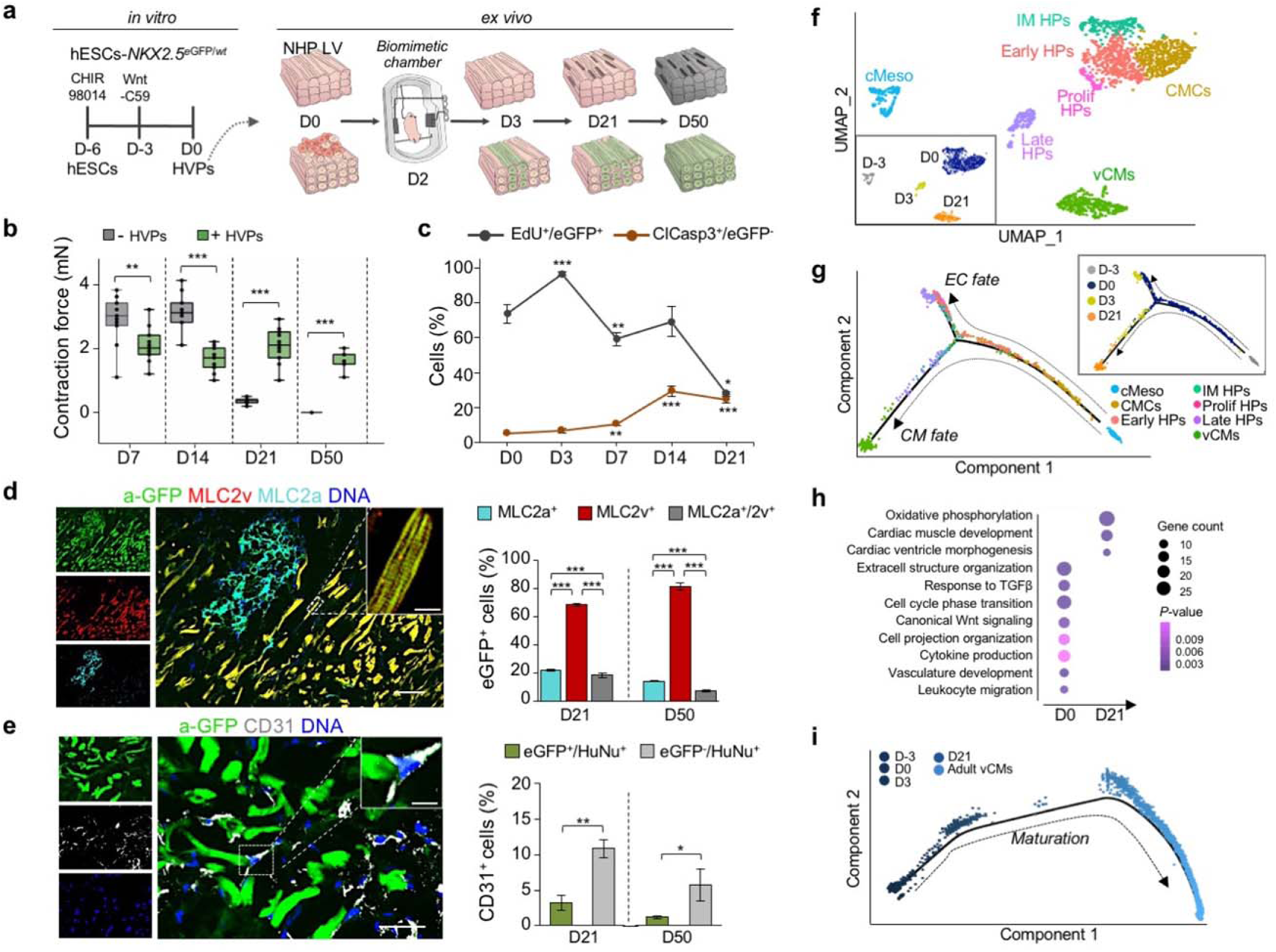
HVPs expand, repopulate and functionally mature in an *ex vivo* 3D NHP heart model. **a**, Schematic of the experimental setup for *in vitro* differentiation of HVPs from *NKX2-5*^eGFP/*wt*^ hESCs (left) and their *ex vivo* co-culture with native NHP LV slices in biomimetic chambers (right). **b**, Contractile force of *ex vivo* cultured NHP heart slices with and without HVPs on indicated days (D) of co-culture. Box plot shows all data points as well as the median and quartiles; n = 11 patches/time point; **p<0.005, ***p<0.001 (*t*-test). **c**, Percentage of EdU^+^/eGFP^+^ and ClCasp3^+^/eGFP^-^ cells during co-culture. Data are mean ± SEM; n ≥ 3 samples/time point; *p<0.05, **p<0.005, ***p<0.001 *vs* D0 (*t*-test). **d, e**, Left, representative immunofluorescence images of D50 chimeric human-NHP heart constructs using an antibody against GFP (a-GFP) together with antibodies for MLC2a and MLC2v (**d**) or CD31 (**e**). Scale bar 100 μm in **d**, 50 μm in **e**, 10 μm in insets. Right, percentage of eGFP^+^ cells expressing MLC2v, MLC2a, or both (**d**) and human cells expressing CD31 (**e**) on D21 and D50. HuNu, human nuclear antigen. Data are mean ± SEM; n = 4 samples/time point; **p<0.005, ***p<0.001 (*t*-test). **f**, UMAP clustering of single cells captured on D-3 and D0 of *in vitro* differentiation together with D3 and D21 of *ex vivo* co-culture. cMeso, cardiac mesoderm; CMCs, cardiac mesenchymal cells; Early HPs, early heart progenitors; IM HPs, intermediate heart progenitors; Late HPs, late heart progenitors; vCMs, ventricular cardiomyocytes. **g**, Developmental trajectory analysis of captured cells coloured by population identity and time of collection (inset). EC, endothelial cell. **h**, Representative GO terms upregulated during *ex vivo* co-culture. **i**, Pseudotime trajectory of captured cells combined with adult vCMs from Wang *et al.* 2020. Colour gradient (from dark to light) according to maturation.

To establish a molecular roadmap for HVP specification and maturation during heart repair, we profiled cells on D0 (before seeding on NHP LV slices) and eGFP^+^ cells isolated on D3 and D21 of *ex-vivo* co-culture by single cell RNAseq (scRNAseq). We then integrated the data with our previously published scRNAseq dataset from D-3 of *in-vitro* differentiation^31^. Transcriptomes of over 1,615 cells were recovered. Unsupervised clustering analysis identified 7 distinct sub-populations; stage dependent clustering was evident for all samples (Fig. 1f). On D-3, corresponding to the time point of cardiac lineage commitment^32^, cells expressed high levels of genes typical of early cardiac mesodermal cells, such as *EOMES, MESP1* and *LGR5* (Extended Data Fig. 2a, b and Supplementary Table 1). On D0, cells distributed into 4 distinct clusters: transcriptomes of early (*KRT18*/*ID1*), intermediate (*KRT8*/*PRDX1*), and proliferating (*TOP2A*/*CCNB1-2*) heart progenitor states as well as cardiac mesenchymal cells (*PLCE1*/*PPA1*) were captured (Fig. 1f and Extended Data Fig. 2a, b). Transcripts related to extracellular matrix organization (*DCN*/*TIMP1*/*LUM*/*FN1*/*COL3A1*) and ventricular muscle structure/maturation (*MYL3*/*TTN*/*TNNC1*/*ACTC1*/*PLN*) defined late eGFP^+^ cells and ventricular CMs on D3 and D21 within the myocardial tissue, respectively (Fig. 1f and Extended Data Fig. 2a, b). In order to achieve a temporal resolution of cardiac fate decisions, we aligned cells captured at the various time points in pseudotime, a computational measure of the progress a cell makes along a certain differentiation trajectory^33^. The resulting trajectory began with cardiac mesodermal cells on D-3 followed by mesenchymal cells and early progenitors from D0 and then bifurcated into two lineages where endothelial-committed late progenitors on D3 and HVPs and their derivative CMs on D21 were allocated (Fig. 1g and Extended Data Fig. 2c).

Gene ontology (GO) enrichment analysis of differentially expressed genes (DEGs) in cells from D0 to D21 revealed progressive activation of GO terms related to cardiac ventricular morphogenesis and maturation, while signalling pathways relevant to the cardiac progenitor state, such as extracellular matrix organization, cell cycle, and canonical Wnt signalling, were gradually suppressed (Fig. 1h). Interestingly, genes upregulated in progenitors at the early time of co-culture also associated with cell migration, cell projection organization, cytokine production, and response to TGFβ (Fig. 1h), suggesting a specific sensing-reacting response of HVPs to the tissue environment. Of note, vasculature development was likewise enriched, confirming the additional potential of some early precursors to differentiate into vessels. To better define the level of maturation achieved by the HVP-derived CMs, we integrated our data with a published scRNAseq dataset of *in vivo* human adult ventricular muscle^34^ in pseudotime (Fig. 1i and Extended Data Fig. 2d). eGFP^+^ cells on D21 partially allocated together with adult ventricular CMs at the end of the differentiation trajectory and expressed high levels of structural, functional, and metabolic genes characteristic of the adult state (Fig. 1i and Extended Data Fig. 2d). Taken together, our single cell transcriptomic analyses allowed the construction of a differentiation route through which early mesodermal cardiac progenitors generate mature CMs in response to the signalling cues of a gradually dying myocardium.

## HVPs directly migrate to and remuscularize the damaged myocardium in an acute NHP heart injury model

Embarking on the heart tissue slices *ex vivo*, we designed a model of acute cardiac injury to provoke tissue death and elucidate HVP properties in response to injury signals (Fig. 2 and Extended Data Fig. 3). Radiofrequency ablation (RFA) is routinely employed in patients with atrial fibrillation to terminate arrhythmogenic foci. We established 20W for 15sec as efficient RFA conditions to destroy, in a standardized manner, a defined area of the cellular compartment within the NHP heart slices, leaving the extracellular matrix (ECM) as scaffold intact (Extended Data Fig. 3a). A progressive invasion of activated cardiac fibroblasts (CFs) expressing the discoidin domain receptor 2 (DDR2) as well as an increase of collagen type I deposition in the RFA-injured area were visible over time, with a complete scarring of the tissue by D21 (Extended Data Fig. 3b). In the first series of experiments, we seeded equal amounts of *NKX2.5*^eGFP/*wt*^ HVPs or CMs onto bio-printed pluronic frames on one side of the NHP heart slices, generated RFA injury on the opposite side, and evaluated the cellular response to the damage by live cell imaging of the eGFP signal (Fig. 2a). In contrast to CMs, HVPs departed from their local seeding site and migrated in a directed manner towards the injured region, colonising it within 4 days (Fig. 2a and Extended Data Fig. 3c, d). By D15, HVPs had differentiated into CMs and the RFA area appeared largely remuscularized, with new eGFP^+^ CMs being elongated and showing aligned myofibrils with organized sarcomeres on D21 (Fig. 2b). A significant reduction of scar volume was measured exclusively in HVP-treated heart slices and, consistently, contractile function of the tissue was considerably improved (Fig. 2c, d and Extended Data Fig. 3e). To assess the potential of HVP-derived CMs to functionally integrate into the electromechanical syncytium after injury, we performed real-time intracellular Ca^2+^ analysis comparing regions of interest (ROI) within the damaged and native myocardium. In contrast to CM-treated heart slices, Fluo-4 fluorescence clearly propagated through the injury area when HVPs had been applied, with differentiated HVPs displaying [Ca^2+^]i oscillations similar to and synchronized with those in adjacent native NHP CMs (Fig. 2e), indicating electromechanical integration of the HVP-derived CMs.

**Figure 2.**
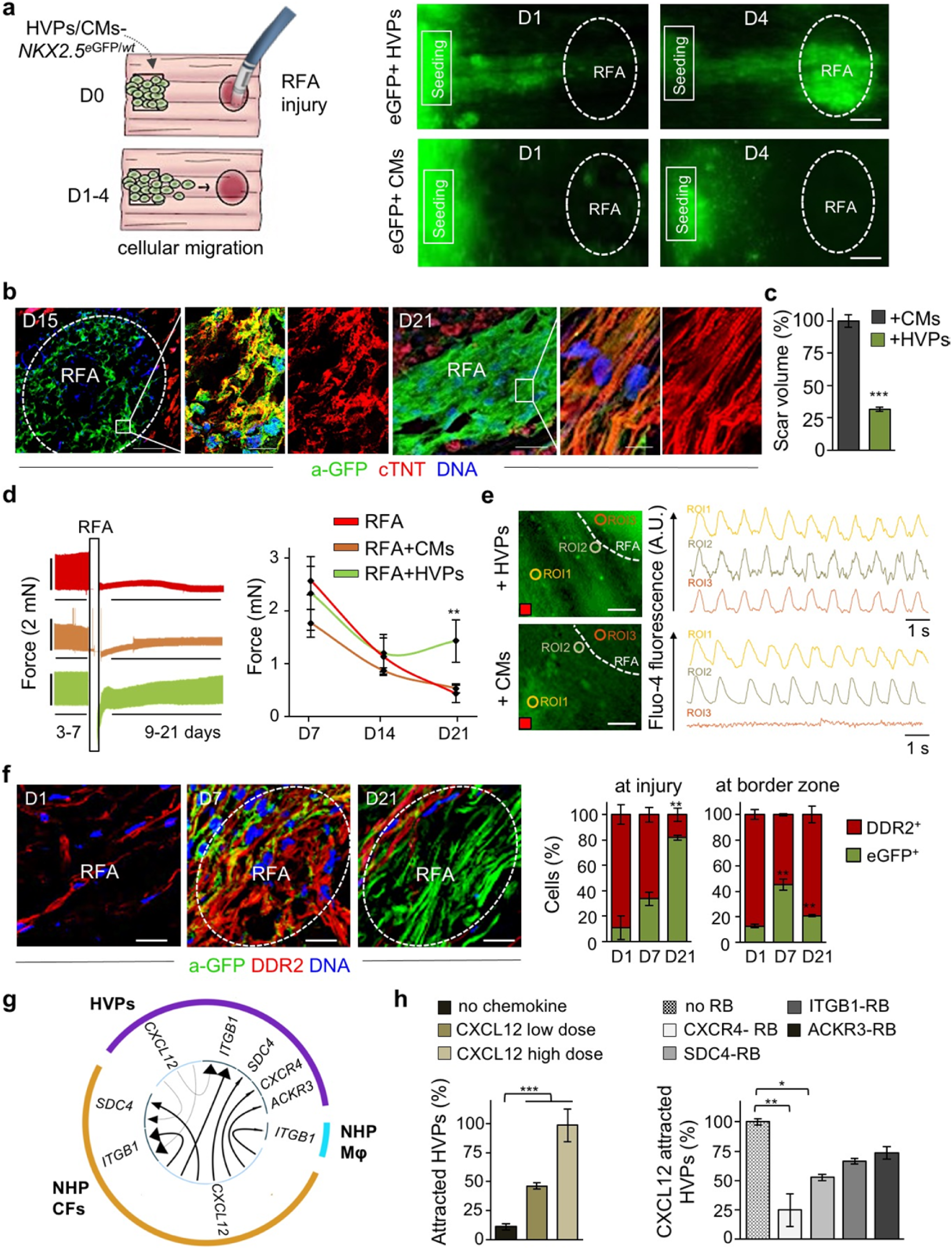
HVPs are chemoattracted to sites of cardiac injury *via* CXCL12/CXCR4 signalling and remuscularize the scar. **a**, Left, schematic of experimental design for selective seeding of *NKX2-5*^eGFP/*wt*^ hESC-derived HVPs or CMs onto bioprinted pluronic frame on NHP heart slices and standardized radiofrequency-ablation (RFA) injury on the opposite tissue site. Right, sequential live-imaging of eGFP signal at indicated days. Scale bars 200 μm. **b**, Representative immunostaining of eGFP and cardiac troponin T (cTNT) in NHP constructs on D15 and D21 after RFA. Magnifications of the framed areas are shown in adjacent panels. Scale bar 200 μm for D15, 100 μm for D21, 10 μm for magnifications. **c**, Statistical analysis of relative reduction of scar volume with HVPs compared to CMs on D21. n=2 patches per group, ≥ 28 z-stack images/patch. **d**, Left, Representative recordings of contractile force before and after RFA, separated by a blanking period of 2 days for re-adjustment of preload (left) and corresponding statistical analysis (right). n=3 samples/condition. **e**, Representative images of Fluo-4 loaded NHP-HVP and -CM constructs (left) and corresponding Ca^2+^ transients at indicated regions of interest (ROI) (right). Scale bar 100 μm. Red box indicate stimulation point (1Hz). **f**, Left, representative immunostaining of eGFP and DDR2 in NHP constructs at indicated days after RFA. Scale bars 200 μm. Right, percentage of eGFP^+^ and DDR2^+^ cells at RFA injury or border zone; n=3 samples/time point. **g**, Circos plot for ligand-receptor pairing showing top ten interactions identified in scRNAseq of NHP-HVP constructs at 24 and 48 hours after RFA injury and HVP application. Fraction of expressing cells and link direction (chemokine to receptor) are indicated. Mφ, macrophages. **h**, Percentage of chemoattracted HVPs in trans-well migration assays in absence and presence of low (20 ng/ml) or high (80 ng/ml) dose of CXCL12 (left) or after addition of the indicated receptor blockers (right). n=3 samples/condition. All data are indicated as mean ± SEM; **p<0.05, **p<0.005, ***p<0.001 (*t*-test).

To further dissect the cellular and molecular mechanisms underlying the observed HVP directed migration towards the RFA injured tissue and the subsequent positive remodelling during the scarring process, we first evaluated the dynamic cellular composition of the tissue around and at the injury site over time. Immunofluorescence analysis indicated that, one day after RFA, activated DDR2^+^ NHP CFs were heavily populating the border zone and already reached the damaged area before the human eGFP^+^ HVPs; both cells coexisted in the injured and surrounding regions after 1 week (Fig. 2f). Shortly after, the RFA site was predominantly colonized by human eGFP^+^ cells and the border zone from NHP DDR2^+^ CFs (Fig. 2f). These observations suggested that cell-cell communication through chemokines or physical interaction between the host CFs and the human progenitors might instruct HVP migration, differentiation, and scar remodelling.

## Chemotaxis of HVPs to sites of cardiac injury is mediated by CXCL12/CXCR4 signalling

During development ISL1^+^ HVPs are highly migratory for heart tube extension and outflow tract formation^23^. To gain insights into the mode of HVP migration in our cardiac injury model, we first performed a trans-well migration assay, where HVPs were placed on top of a permeable membrane and RFA-injured or uninjured NHP heart slices at the bottom (Extended Data Fig. 3f). A significantly boosted migration was observed in presence of RFA. Interestingly, while multiple, homogeneously distributed RFAs prompted HVPs to evenly migrate through the membrane, a directional migration towards the injured area was observed with a single isolated RFA (Extended Data Fig. 3f), indicating the production of a chemoattractant gradient specifically arising from NHP cells at the damaged area.

To further dissect the molecular programs for directed HVP chemotaxis and response, we profiled migrating eGFP^+^ cells (at 24 hours; 485 cells) and arriving eGFP^+^ cells at the RFA injury (at 48 hours; 269 cells) as well as eGFP^-^ tissue resident host cells (315 cells) by scRNAseq (Extended Data Fig. 4a). Cells were embedded in low-dimensional space using UMAP followed by unsupervised clustering. Seven distinct clusters were recovered, which grouped into three main cell populations (Extended Data Fig. 4a, b and Supplementary Table 1). Cluster 1 and 4 belonged to the NHP cell group and mapped to CFs and monocyte/macrophages, respectively. Human cells formed the other 2 major cell groups. One contained 4 clusters, which were classified as early HVPs (expressing high levels of metabolic genes as *MBOAT1, UQCRQ*, and *MT-ND1,2,4,5,6*, but lacking expression of CM transcripts; cluster 0), activated HVPs (*LAMA5, FLRT2*, and *TNC*; cluster 2), proliferating HVPs (*TOP2A, CDC20*, and *CCNB2*; cluster 5), and early ventricular CMs (*MYH6, MYL3, TNNC1*; cluster 6). The second comprised of a homogeneous population of HVPs (cluster 3) characterized by high expression of genes involved in chemotaxis (*NRP1, CCL2-19-21, CXCL2-6-8-12, ITGB1, WASF1, RPS4X, INPPL1*), an unique gene signature not captured before. GO analysis of DEGs between cluster 3 and the other HVP clusters identified enrichment of terms related to cell motility, chemotaxis, actin filament organization, axon guidance cues, and ECM organisation, (Extended Data Fig. 4c), supporting the migratory feature of this cell population. We further directly characterized the intercellular communication signals between HVPs and NHP cardiac cells by performing an *in silico* single cell receptor-ligand pairing screen. We found over-representation of a significant pairing of CXCL12 as ligand with several membrane receptors, including CXCR4, SDC4, ITGB1, and ACKR3 (Fig. 2g). While CXCL12, SDC4, and ITGB1 were expressed in both HVPs and NHP fibroblasts, the CXCR4 and ACKR3 receptors were highly enriched in the HVPs (Extended Data Fig. 4d). Trans-well migration assays under gain- and loss-of-function conditions demonstrated that HVPs exhibited enhanced migratory behaviour under CXCL12 as chemoattractant, which was specifically reduced in presence of blocking antibodies for CXCR4 and SDC4 (Fig. 2h). Notably, binding of CXCL12 to SDC4 is known to facilitate its presentation to the CXCR4 receptor^35^. Collectively, our data support the model that HVPs expressing CXCR4 sense CXCL12 secreted by CFs at the injury site as a chemoattracting signal to repopulate the damaged myocardial compartment. A similar, chemokine-controlled deployment of SHF cells has been identified as intra-organ crosstalk between progenitors and FHF CMs during mouse cardiogenesis^36^, suggesting that migration programs that are functional during cardiac development are re-activated in HVPs during organ regeneration.

## Distinct dynamical cellular states underlie HVP regenerative potential upon tissue injury

To capture all transition cell types and analyse the stepwise process of HVP-mediated cardiac repair, we integrated all scRNAseq data of HVPs on D0, 24h and 48h after RFA injury, as well as HVP-derived CMs on D21 co-culture (2,114 cells) and generated a diffusion map of tissue damage-induced cardiac differentiation (Fig. 3a, b). Heat-mapping of gene expression with cells ordered in the trajectory revealed a temporal sequence of gene expression events and identified cells at intermediate stages of *injury sensing* and *injury response* (Extended Data Fig. 5a). Dot plotting illustrated gene signature shifts among the different stages (Fig. 3c). In the first 24h after injury, HVPs “sense” the tissue damage and activate gene programs for ECM remodelling (e.g. *COL6A1, ADAMTS9, FLRT2*), secretion and response to cytokine (*SPP1, STX8, TGFBI, IL6ST*), as well as initiation of migration (*PLAT*). Subsequently (48h), they upregulate genes typical of migratory cells, including chemoattraction signalling genes (*PLXNA2, CMTM3*, and *CXCL12*), cell motility genes (*SNAI1, SNAI2, FAT1*, and *TIMP1*), and transcripts of cytoskeleton organization (*ARPC2*) as well as axon guidance (*SLIT2, NFIB*, and *UNC5B*) and cell projection (*RGS2, THY1*, and *ITGA1*). In this *migratory* state, gene signatures of secretion (*COPB2, VPS35*, and *SPTBN1*) and cardiac muscle differentiation (*VCAM1, MHY6, PALLD*, and *TMOD1*) become increasingly important as *counteracting* response to injury (Fig. 3c). Indeed, mass spectrometry analysis of supernatants from NHP heart slices 48h after RFA injury revealed a significant upregulation of secreted proteins in the presence of HVPs (Extended Data Fig. 5b). Interestingly, the majority of them are involved in ECM organization (e.g. HSPG2, SPARC, FN1) and fibrotic/inflammation response (e.g. FSTL1, PRDX1, SPTAN1), reinforcing the capability of HVPs to influence scar remodelling.

**Figure 3.**
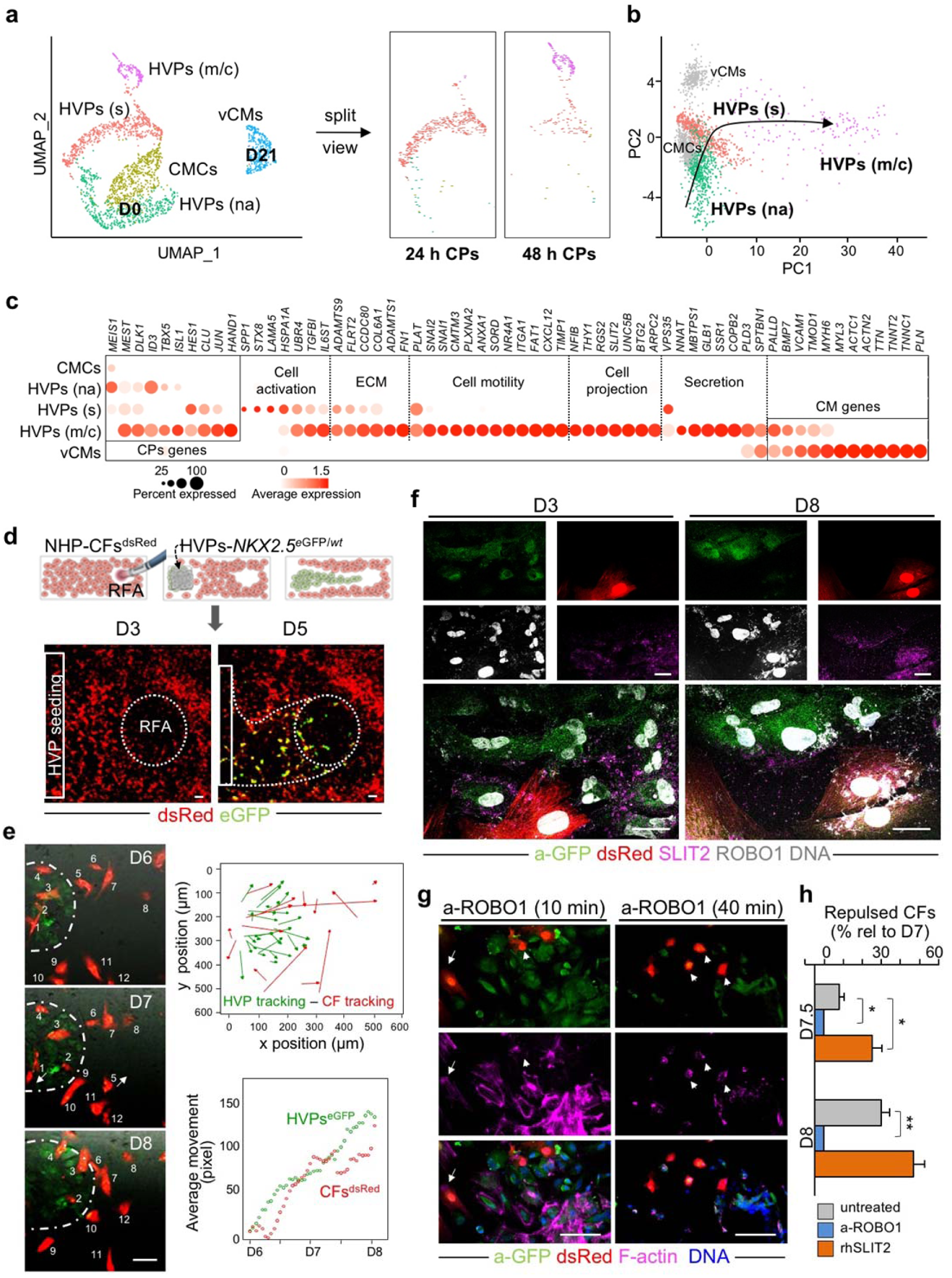
SLIT2/ROBO1 signalling mediates activated CF repulsion and prevents myocardial scarring. **a**, Twenty-four and 48 hours human scRNAseq datasets are integrated with D0 and D21 CM dataset and projected onto UMAP plots, coloured by cluster assignment and annotated post hoc. Both the aligned (left) and split (right) views are shown. CMCs, cardiac mesenchymal cells; HVPs (na), non-activated; HVPs (s), sensing; HVPs (m/c), migrating and counteracting; vCMs, ventricle cardiomyocytes. **b**, PCA plot of different cell clusters, with the principal curve indicating the pathway of injury response. **c**, Dot plot showing gene signature shifts among different dynamic cellular states. The shadings denote average expression and the size of dots the fractional expression. **d**, Top, schematic of 2D model for RFA injury of NHP-CFs expressing dsRed followed by *NKX2-5*^eGFP/*wt*^ HVP seeding and monitoring of co-culture. Bottom, sequential live imaging of dsRed^+^ and eGFP^+^ cells during migration. Scale bars 200 μm. **e**, Left, representative time-lapse images of dsRed^+^ and eGFP^+^ cells at the RFA injury site during CF repulsion on indicated days. Dotted line delineates HVP migration front. Scale bar 100 μm. Right, cell tracking over time (top) and average movement (bottom) analysis of HVPs and CFs. **f**, Representative immunostaining for eGFP, SLIT2, and ROBO1 on D3 and D8. Scale bars 25 μm. **g**, F-actin and eGFP immunofluorescence an D8 after ROBO1 antibody exposure for 10 and 40 minutes. Change of CF shape (arrow head) and F-actin localized on protrusion side of CFs (arrow). Scale bars 75 μm. **h**, Percentage of repulsed CFs at the injured site analyzed on D7.5 and D8 in standard condition (untreated) or after ROBO1 antibody and rhSLIT2 treatment on D7. Data are normalized to D7 and presented as mean ± SEM, n=3. \**p*<0.05, \*\**p*<0.005 *vs* untreated (*t*-test).

## SLIT2/ROBO1 mediates activated fibroblast repulsion and reduces scar formation

CFs play an essential role in heart development and repair. During heart regeneration in zebrafish, activation of resident fibroblasts support CM growth and maturation through secretion of specific ECM components^2,37^. To further investigate the temporal and spatial direct crosstalk between CFs and HVPs in our *ex vivo* cardiac injury model, we isolated CFs from native NHP hearts, stably expressed dsRed by lentiviral transduction, and performed live imaging during monolayer co-culture with *NKX2.5*^eGFP/*wt*^ HVPs. RFA injury (20W for 7sec) was performed on one site of the dsRed^+^ CF monolayer and seeding of *NKX2.5*^eGFP/*wt*^ HVPs on the other (Fig. 3d). Similar to the native tissue, the first cells invading the injured area were dsRed^+^ CFs, followed by eGFP^+^ HVPs within 5 days (Fig. 3d). Remarkably, while HVPs were directly chemoattracted to the injury, CFs appeared dynamically repelled at the contact sites with migrating HVPs (Fig. 3e and Extended Data Movie 1). Live cell tracking of more than 100 cells over 3 days demonstrated that the majority of CFs, after interacting with HVPs, indeed moved actively away from the HVP migratory path and were repelled from the injured area when the HVPs started to densely populate it on D8 (Fig. 3e). Immunocytochemistry of filamentous (F)-actin revealed a specific retraction of cell protrusions precisely occurring at cellular contact sites with the HVPs (Extended Data Fig. 6a), suggesting that the latter possibly control actin dynamics of CFs at the interaction sites. Given the detected upregulation of genes involved in axon guidance in the migratory HVP state, including SLIT2, we postulated that SLIT2/ROBO1 signalling, a known repulsive guidance cues for neuronal axons^38^, might control HVP-mediated CF repulsion by regulating cytoskeletal organization and cell motion. Co-immunofluorescence analysis demonstrated expression of both SLIT2 ligand and ROBO1 receptor in migrating HVPs on D3, while no signal was detected in the surrounding CFs (Fig. 3f). On D8, however, co-localization of SLIT2 and ROBO1 was observed mainly at the membrane of repulsed CFs, with enriched SLIT2 signal at the contact sites with the HVPs (Fig. 3f). Quantitative RT-PCR confirmed that SLIT2 was produced by the HVPs and ROBO1 was expressed in both cell types at the stage of CF repulsion (Extended Data Fig. 6b). Loss-of-function experiments using an antibody blocking ROBO1 signalling substantiated that, under ROBO1 inhibition, HVPs failed to induce actin polymerization and lamellipodia formation in the interacting CFs, leading to reduced CF motility and lack of repulsion (Fig. 3g, h and Extended Data Fig. 6c). No effects were observed in the distant CFs (Extended Dada Fig. 6c). Conversely, treatment with recombinant human SLIT2 enhanced F-actin content and membrane protrusions specifically in CFs communicating with HVPs (Extended Data Fig. 6d), resulting in enhanced repulsion (Fig. 3h).

## HVPs regenerate injured porcine myocardium *in vivo* without inducing arrhythmias

To investigate the full regenerative potential of HVPs, we performed allogeneic *in vivo* transplantation experiments in genetic modified pigs ubiquitously expressing LEA29Y, a human CTLA4-Ig derivative that blunts systemic T cell response^39^. This immuno-compromised model offers an ideal setting for testing human cell therapies in xenotransplantation approaches, enabling improved graft survival. Two epicardial RFA injuries (25W for 7sec) were induced afar in the anterior heart wall and 6×10^7^ *NKX2.5*^eGFP/*wt*^ HVPs were injected ∼1cm apart from one damaged site, while the other served as control (Fig. 4a and Extended Data Movie 2). Morphological assessment of RFA-induced tissue damage in freshly isolated wild-type porcine hearts demonstrated consistent size of myocardial injury (Fig. 4b, c). LEA29Y animals were treated daily with methylprednisolone and tacrolimus and euthanized under full anaesthesia on D3 (n=1), D5 (n=2), and D14 (n=2) post-transplantation. None showed any macroscopic signs of tumour formation (Extended Data Fig. 7a). Immunohistology on D3 documented a directed, guided migration of eGFP^+^ HVPs towards the RFA-injured area (Fig. 4d). On D5, eGFP^+^ cells reached the RFA site, appeared mainly in clusters, and repopulated 6.3±0.6% of the scar (Fig. 4d, f). No significant difference in scar volume was measured at this time point between HVP-populated and control scars (Fig. 4e). However, 2 weeks after injury, eGFP^+^ cells constituted 21.0±2.9% of the injured area and scar volume was significantly reduced to half compared to control scars (Fig. 4d, e). Remarkably, their spatial distribution throughout the scar inversely correlated with the depth of the analysed plane, denoting the highest concentration of eGFP^+^ cells at the more epicardial layers with the largest damaged area (Fig. 4f). Immunofluorescence analysis demonstrated that the vast majority of eGFP^+^ cells engrafted in the damaged tissue were cTNT^+^ CM with elongated shape, aligned myofibrils, and well-organized sarcomeres (Fig. 4g). Expression of the gap junction protein connexin 43 was detected at the intercalated discs of eGFP^+^ CMs and at the contact zone between graft and host CMs (Fig. 4g). Moreover, immunostaining for the endothelial marker CD31 documented significantly enhanced neo-angiogenesis at the RFA site after HVP transplantation (Fig. 4h and Extended Data Fig. 7b). Notably, ∼6% of CD31^+^ cells were of human origin (Fig. 4h). No evidence of acute graft rejection in the transgenic LEA29Y pigs under the immunosuppressive regimen was observed on D14 post-transplantation, as assessed by CD68 immunodetection. Interestingly, we even observed a reduction of CD68^+^ cells at both injured and adjacent areas in HVP-treated RFA compared to control (Extended Data Fig. 7c), suggesting that HVPs might directly influence post-injury inflammation.

**Figure 4.**
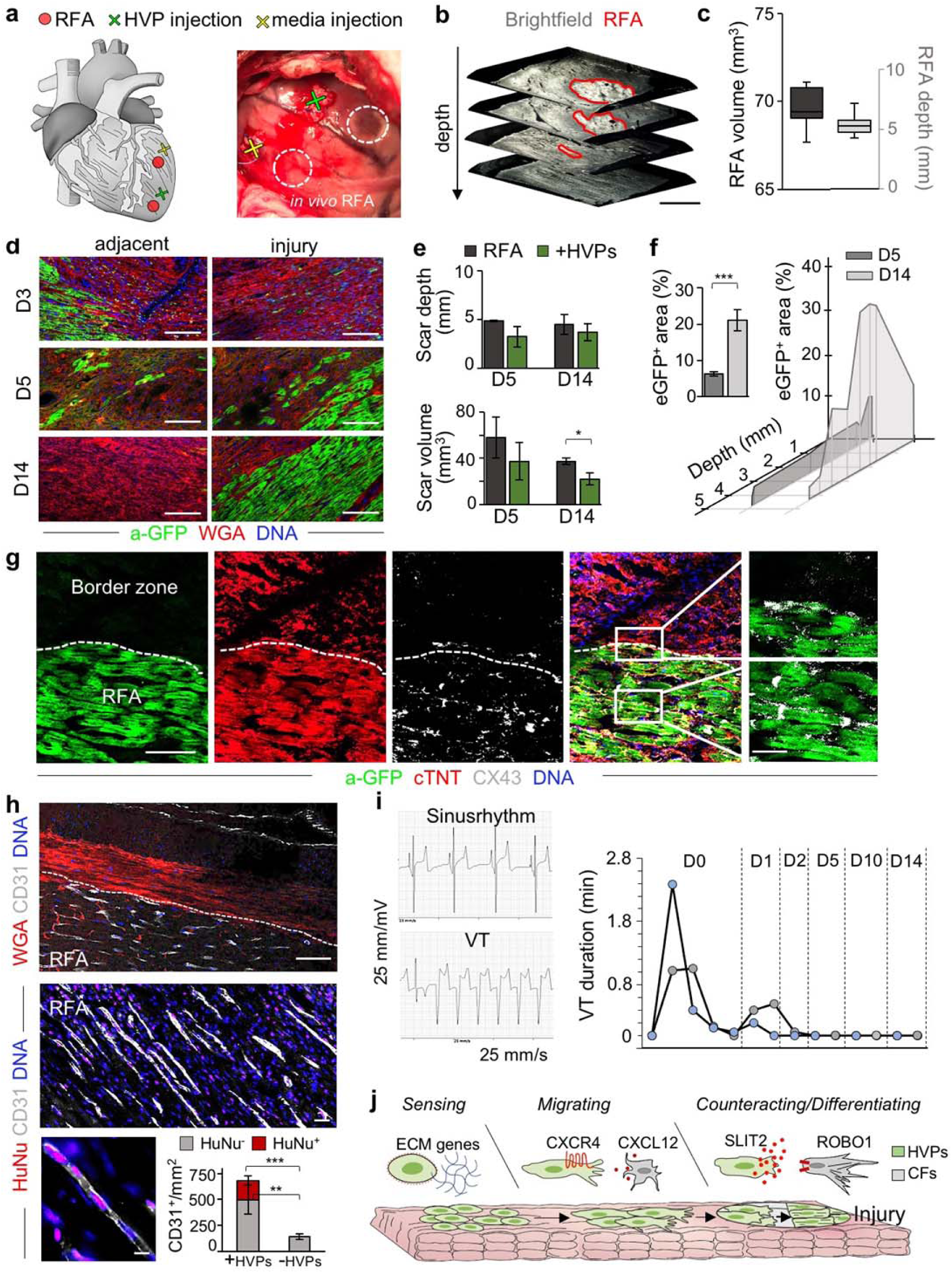
HVPs directly migrate towards the damaged myocardium and remuscularize the injured tissue without arrhythmias in a transgenic LEA29Y porcine model *in vivo*. **a**, Schematic of *in vivo* experimental design depicting 2 left ventricular RFA injuries and adjacent injection of HVPs or cell-free media. **b, c**, Representative 3D reconstruction of non-transmural RFA injury (**b**, scale bar 2 mm) and statistical analysis of scar volume and depth of RFA injuries in freshly explanted wild-type pig hearts indicating standardized injury size (**c**). The median and quartiles are shown; n=3. **d**, Representative fluorescence images of injury and adjacent sites after WGA and anti-GFP (a-GFP) co-staining on D3, D5 and D14. Scale bars 100 μm. **e**, Quantification of *in vivo* scar depth and volume on D5 and D14 with or without HVP injections. **f**, Percentage of eGFP^+^ area within the RFA injury (left) and according to depth of the cutting plane. **g**, Representative immunostaining of eGFP, cTNT, and CX43 in RFA and border zone on D14. Magnifications on the right correspond to the boxed area in the merge image. Scale bar 50 μm and 10 μm (magnifications). **h**, Representative fluorescence images of HVP-treated RFA injury site after immunostaining for CD31 in combination with WGA (top) or with anti-human nuclei (HuNu, middle and bottom). Scale bar 50 μm top, 25 μm middle, 10 μm bottom. Bar graph shows the average number of CD31^+^ cells/mm^2^ cells derived from host (HuNu^-^) or human progenitors (HuNu^+^) in HVP-treated and untreated RFAs. **i**, Eventrecorder readout with representative ECG traces of sinus rhythm (top) and degeneration to ventricular tachycardia (VT, bottom). Right, occurrence and duration of VTs at the indicated days after transplantation. n=2 eventrecorder traces. **j**. Scheme depicting the identified dynamical cellular states of human HVPs during tissue heart repair and the involved signalling pathways. Data in **e, f**, and **h** are mean ± SEM, n=2 per group. * p<0.05, **p<0.005, ***p<0.001 (*t*-test).

Ventricular arrhythmias have emerged as major side effect of CM cell therapy. Electrophysiological studies in large animals consistently observed arrhythmias originating in graft regions from ectopic pacemakers^40,41^. To assess the electrophysiological consequences of hPSC derived HVPs, we subjected the two pigs euthanized on D14 to permanent ECG monitoring following cell therapy and RFA injury using an implantable event recorder. No ventricular tachycardia (VT) was observed before cell transplantation and only few non-sustained VT episodes occurred shortly within the first 48h of treatment (Fig. 4i).

## Discussion and future directions

Our *ex vivo* chimeric model of human HVPs and NHP heart tissue provides an unprecedented system to refine molecular pathways implicated in cardiac regeneration and healing at a single cell resolution. As such it offers an innovative approach to predict outcome of cell-based regeneration with high fidelity, which could be applied to other non-regenerative organs such as brain. We demonstrate that HVPs harbour the unique potential to sense and counter-act injury by reactivating sequential developmental programs for directed migration, fibroblast repulsion, and ultimate muscle differentiation in the scar area (Fig. 4j).

scRNAseq data unravelled key signalling pathways underlying HVP-mediated heart repair and scarless healing during an acute injury response, including SLIT2/ROBO1. It will be of particular interest to evaluate whether pharmacological manipulation of such signalling pathway could circumvent cell application. Moreover, future studies should assess whether HVP therapy could be beneficial in clinical settings of chronic heart failure (e.g. ischemic heart disease, genetic cardiomyopathies, and post-myocarditis) to reduce pre-existing fibrosis. Recently, the ESCORT trial performed first transplants of hPSC-derived cardiac progenitors surgically delivered as patches onto the heart’s surface in patients with ischemic cardiomyopathy and reported no adverse side effects^18^. In parallel, translational efforts have evaluated the potential of hPSC-derived CMs to engraft in the heart of large animals, including primates. Electromechanical coupling and improvement of systolic LV function have been reported^17,40,41^. However, concerns remain regarding the immaturity of engrafted CMs, survival of the cells and the propensity for ventricular arrhythmias. We found that *in vivo* hPSC-derived HVPs transplanted in the injured myocardium of LEA29Y transgenic pigs did not induce sustained VTs over a two week period. Yet, arrhythmogenic potential of cell grafts needs to be further assessed in long-term analyses. We also observed an increased neovascularization *in vivo*, with a proportion of endothelial cells being of human origin. Further studies need to demonstrate whether such neovascularization response will be sufficient to restore normal blood flow. Robust arterial input will be crucial for permanent functional improvement, which may require a combination of cell therapy with other modalities.

In conclusion, our data indicate that HVPs harbour the unique capability to target both loss of myocardium and fibrotic scarring in the primate heart, and support their therapeutic potential. However, before HVP transplantation can be translated to humans much work remains to determine whether pharmaceutical-grade batches of HVPs (purity ≥90%, yields ≥10×10^7^ cells) can be achieved, safety risks related to ventricular arrhythmias can be excluded, and the use of hypo-immunogenic PSC lines can circumvent long-term rejection. Developing innovative therapeutic strategies that are rooted in fundamental biology of cardiac development could pave the way for successful cell-based cures of heart disease.

## Methods

### ESC maintenance, cardiac differentiation, and HVP MACS-based purification

Embryonic stem cell lines ES03 *NKX2.5*^eGFP/*wt*^ and H9 *NKX2.5*^eGFP/*wt* 42^ were generously gifted from Dr. David Elliott, MCRI Australia) and maintained on Matrigel-coated plates (BD Biosciences, Germany) in E8 medium (Gibco, USA) with daily medium change. After reaching a confluency of 85-90%, cells were passaged by dissociation into single cells using Accutase (Innovative Cell Technologies, USA) at 37°C for 5 minutes and replated in a ratio of 1:6 or 1:9 on new Matrigel-coated plates in E8 supplemented with 5 μM ROCK inhibitor Y-27632 (Stemcell Technologies, Canada) for 24h. Differentiation to HVPs was achieved according to our previously published protocol^28^. Briefly, after dissociation with Accutase, ESCs were plated on Matrigel-coated cell culture dishes at a density of 1×10^6^/well in E8 supplemented with 5 μM ROCK for 24 hours followed by culture in E8. When full confluency was reached, differentiation was initiated on day -6 by adding RPMI/B27 minus insulin (Gibco, USA) supplemented with 1 μM CHIR 98014 (Selleckchem, USA). Media was changed to RPMI/B27 minus insulin (HVP culture medium; CCM) after 24 hours. On day -3, a combined medium consisting of 1 ml collected conditioned media and 1 ml fresh CCM supplemented with 2 μM Wnt-C59 (Selleckchem, USA) was applied and completely replaced by CCM on day -1. On day 0 HVPs were collected for MAC sorting. After dissociation into single cells with Accutase, cells were stained with Anti-TRA-1-60 MicroBeads (Miltenyi Biotec, Germany) before negative sorting with an autoMACS Pro Separator (Miltenyi Biotec, Germany) according to manufacturer’s instructions. A fraction of the cells was stained with an anti-human TRA-1-60 antibody conjugated with Alexa Fluor 488 (StemCell Technologies, Canada) and analysed by flow cytometry using a BD FACSCantoII according to manufacturer’s instructions. Batches with <5% TRA-1-60 positive cells were used for *in vivo* transplantation experiments. To generate mature CMs, HVPs were cultured further in 12 well plates in RPMI/B27 containing insulin and medium was changed every other day. For long-term storage, cardiac progenitors (D0) or differentiated cardiomyocytes (D25) were frozen in CryoStor cell cryopreservation media (Sigma Aldrich, USA).

### *Ex vivo* NHP heart slice culture

For *ex vivo* heart slice cultivation, freshly explanted NHP left-ventricular myocardial tissue was placed in 2,3-Butanedione 2-Monoxime (BDM; Sigma Aldrich, USA) at 4°C and shipped from the German primate centre, Göttingen, where the primate had been euthanized in course of a vaccination study (file reference: 33.19-42502-04-16/2264), from Karolinska Institutet, Sweden (file reference: N 277/14) or from the Walter Brendel Institute, LMU, Germany (file reference: ROB-55.2--2532.Vet_02-14-184). Within 12-24 hours, heart tissues were sectioned on vibratome (VT1200S, Leica Biosystems, Germany) to approximately 1 cm x 2 cm x 300 μm thick tissue slices. Slices were anchored in biomimetic cultivation chambers (BMCC) via small plastic triangles attached to the slices with tissue adhesive (Histoacryl; B. Braun, Germany) according to fiber direction and subjected to physiological preload of 1 mN and stimulation at 1 Hz (50 mA pulse current, 1 ms pulse duration), as previously described^29^. The slices were maintained in M199 medium (Sigma Aldrich, USA) supplemented with 1% Penicillin-Streptomycin, 0.5% Insulin/Transferrin/Selenium (Gibco, USA) and 50 μM 2-Mercaptoethanol. Medium was replaced every other day (2/3 fresh medium, 1/3 conditioned medium). The BMCCs were anchored on a rocker plate, placed in an incubator at 37°C, 5% CO_2_, 20% O_2_ and 80% humidity. Contractile force of the constructs was continuously measured and contractility data were imported into and analysed by LabChart Reader software (AD Instruments, New Zealand).

### Generation of chimeric human-NHP heart constructs

Native NHP heart slices within BMCCs were underpinned with a hand-trimmed filter (0.40 μm pore size) and suspended in 0.5 ml CCM underneath the filter. For homogenous cell seeding, 2×10^6^ HVPs were thawed and immediately seeded onto the tissue within a pluronic F-127 (concentration of 0.33 g/ml, Sigma Aldrich, USA) frame using a bioprinting device (CANTER Bioprinter V4, Germany), equipped with a 0.58 mm standard Luer-Lock nozzle (Vieweg^®^ Dosiertechnik, Germany). For selective cell seeding, 0.5×10^6^ HVPs or D25 CMs were used. In the first 12 hours after seeding, chimeric constructs were cultured in 500 μl CCM supplemented with 5 μM ROCK inhibitor underneath the filter within BMCCs without rocking and electrical pacing. Twelve-24 hours after seeding, another 500 μl CCM supplemented with 5 μM ROCK inhibitor were added under the filter and rocking (60 rpm, 15° tilt angle) was resumed. After 24h, medium was entirely replaced with 1 ml CCM. On day 2, co-culture slices were maintained in 2.4 ml CCM with continuous electrical pacing (1 Hz) and rocking. Medium was replaced every other day (2/3 fresh medium, 1/3 condition medium).

### RFA-induced myocardial injury

Native NHP heart slices with selectively seeded HVPs or CMs were injured on the opposite tissue border 3 days after cell seeding by applying 20 W for 15 sec using a THERMOCOOL^®^ SF Uni-Directional Catheter, tip electrode 3.5 mm (Biosense Webster, USA) and Stockert 70 radiofrequency (RF) generator (Biosense Webster, USA). During the RFA procedure, physiologic preload was reduced to 0.5 mN within BMCC that was readjusted to 1 mN after 2 days. For *in vivo* experiments, epicardial radiofrequency ablation with 25 W for 7 sec was performed to produce a standardized, non-transmural injury.

### Proliferation analysis by flow cytometry

For quantification of proliferation, HVPs on day 0 of monolayer culture and day 3, 7 and 14 of co-culture were incubated with 10 μM EdU for 24 hours, dissociated with 480 U/ml collagenase type II, fixed with 4% paraformaldehyde (PFA) for 15 min at room temperature (RT), washed three times with PBS and processed using the Click-iT EdU594 Flow Cytometry Assay Kit (Thermo Fisher, USA) according to manufacturer’s instructions. Flow cytometry data were acquired with a Gallios flow cytometer (Beckman Coulter, USA) and evaluated with Kaluza software version 1.2 (Beckman Coulter, USA).

### Ca^2+^ imaging of RFA-injured heart slices

NHP heart slices seeded with HVPs/CMs after RFA injury were loaded with 3 μM Fluo-4-AM in CCM (without phenol red) supplemented with 0.75% Kolliphor EL (Sigma Aldrich, USA) by incubation at 37°C for 60 minutes, washed, and incubated for another 30 min at 37°C to allow de-esterification of the dye in Tyrode’s solution supplemented with Ca^2+^ (135 mM NaCl, 5.4 mM KCl, 1 mM MgCl_2_, 10 mM glucose, 1.8 mM CaCl_2_, and 10 mM HEPES; pH7.35). Calcium signals from native NHP CMs and seeded HVPs were subsequently imaged using an inverted microscope DMI6000 B (Leica, Germany) equipped with a 10x objective, a GFP filter set and a Zyla V sCMOS camera (Andor Technology, UK). Point stimulation electrodes were connected to an HSE stimulus generator (Hugo Sachs Elektronik, Germany) providing depolarizing pulses (40 V, 3 ms duration) at 1 Hz on the tissue border opposite the RFA injury. Imaging settings (illumination intensity, camera gain, binning) were adjusted to achieve an optimal signal to noise ratio with an imaging rate of 14 Hz. ImageJ ROI Manager was used to quantify fluorescence over single cells and background regions. Subsequent analysis was performed in RStudio using custom-written R scripts. After subtraction of background fluorescence, the time course of Fluo-4 signal intensity was expressed as arbitrary units.

### Cell isolation for scRNA-sequencing

For scRNAseq, co-culture patches on day 0, 3 and 21 without RFA, as well as on day 1 and 2 after RFA were dissociated using 20 U/ml papain^43^ (Worthington Biochemical, USA), filtered through a 70 μm filter and resuspended in 3% BSA in PBS. Using FACSAria III (Becton Dickinson), eGFP^+^ cells on day 0, 3 and 21 without RFA and eGFP^+^ and eGFP^-^ cells on day 1 and 2 after RFA were sorted into individual wells of 384-well plates containing Smart-Seq2 cell lysis buffer (ERCC 1:4 ×10^7^ dilution). The lysed single cells were stored at -80°C prior to complementary deoxyribonucleic acid (cDNA) synthesis, using the Smart-Seq2 protocol. The quality of the cDNA was confirmed using Agilent Bioanalyzer (Agilent, USA) and RNAseq libraries were prepared using in-house compatible Tn5 and Nextera index primers (Illumina, USA). Following a final clean-up, the size distribution of the sequence libraries was verified using Agilent high-sensitivity chip and the concentration of each library was measured using the Qubit3 Fluorometer (Thermo Fisher, USA).

### scRNAseq and gene expression analysis

scRNAseq was performed at the sequencing facility of the Karolinska Institutet using the Genome Analyzer HiSeq2500 (Illumina, USA) for single-end sequencing of 56 bp. The Genome Analyzer Analysis Pipeline (Illumina, USA) was used to process the sequencing files of raw reads in the FASTQ format. The cDNA insert was aligned to the hg19/Mmul_1 reference genomes using Tophat2, combined with Bowtie2. Only confidently mapped and non-PCR duplicates were used to generate the gene-barcode cell matrix. Further, quality check steps, including the identification of highly variable genes, dimensionality reduction, standard unsupervised clustering algorithms and the differentially expressed genes analysis were performed using the standard Seurat R pipeline^44^.

### Cell clustering, UMAP visualization and marker-gene identification

The gene-barcode matrix was scaled, normalized and log-transformed. The dimensionality of the data was reduced by principal component analysis (PCA) (20 components) first and then with UMAP (resolution = 0.3). Then, all cells from each cluster were sampled and differentially expressed genes across different clusters were identified with the FindAllMarkers and FindMarkers functions of Seurat R package. Clusters were assigned to known cell types on the basis of cluster-specific markers (Supplementary Table 1).

### Integrated analysis of single-cell datasets

To integrate and validate the robustness of our analysis, we took advantage of two recently published single cell profiles^31,34^. To reduce batch-effect differences, we used the Seurat alignment re-scaling and re-normalizing for the integrated dataset. For all new integrated datasets, we identified variable genes generating a new dimensional reduction that was used for further analysis. Pseudotemporal ordering was done using Monocle 2^45^. In brief, an integrated gene-expression matrix was constructed as described above. With the function differentialGeneTest we analysed differentially expressed genes across different development conditions. At max the top 3,000 genes with the lowest q value were used to construct the pseudotime trajectory.

### Determination of biological processes and molecular function on the basis of enrichment analysis

Statistical analysis and visualization of gene sets were performed using the clusterProfiler R package^46^. GSEA databases were used to determine the enrichment of biological processes, cellular components and molecular function on the basis of the genes that were significantly upregulated. Process-specific signatures were defined by the top genes as ranked by the significance and expression scores.

### NHP CF isolation and lentiviral transduction

Freshly explanted NHP left ventricular myocardium was minced into small pieces and incubated with 550 U/ml collagenase II (Worthington, USA) at 37°C for 15 minutes in a series of 6 digestions. Every 15 minutes the supernatant was collected and centrifuged at 300 g for 5 minutes and washed twice with DMEM/F-12 (Gibco, USA). Isolated cardiac fibroblasts (CFs) were cultured in CF Medium (CFM; DMEM-F12, 10% fetal bovine serum, 2 mM L-Glutamine, 0.5% Penicillin/Streptomycin) at 37°C and 5% CO_2_ with media change every other day. For CF passaging 0.05% Trypsin-EDTA (Gibco, USA) was used.

For lentiviral transduction, a dsRed-expressing lentivirus was produced using a pRRLsin-18-PGK-d transfer plasmid combined with the packaging plasmid (pCMVdR8.74) and the envelope plasmid (pMD2.VSV.G) in HEK293T cells. The CFs were incubated with PGK-dsRed lentivirus and 8 μg/mL of polybrene hexadimethrine bromide (Sigma Aldrich, USA) for 24 hours at 37°C, 5% CO_2_ and the transduction efficiency was evaluated by dsRed expression after 96 hours.

### Monolayer co-culture of HVPs and CFs for cell interaction studies after RFA injury

NHP-CF^dsRed^ (1×10^4^/well) were seeded in 4-well chamber slides (Thermo Fisher, USA) coated with fibronectin (Sigma Aldrich, USA). After 3 days, RFA injury (20 W, 7 sec) was introduced on one border of the chamber slide followed by seeding of 5×10^5^ HVPs on the opposite side. Cellular migration and interaction were studied by time-lapse microscopy (image acquisition every 90 minutes for 3 days; daily medium change with CCM). For the analysis of CF repulsion, anti-ROBO1 (5 μg/ml) and rhSLIT2 (2 μg/ml) treatments (Supplementary Table 2) were performed on days 7 and 8 after RFA injury and HVP seeding and stopped after 10 minutes, 40 minutes and 24 hours. Video clips were analysed with ImageJ for cell movement of HVPs and CFs by TrackMate plug-in^47^.

### Identification of migratory signaling by trans-well migration assay

For trans-well migration studies, the CytoSelectTM Cell Migration and Invasion Assay (Cell Biolabs, USA) was used and samples were processed according to manufacturer’s instructions. In brief, 0.5×10^6^ HVPs (D0) were suspended in 300 μl serum-free medium and plated on the upper compartment of the trans-well migration assay (polycarbonate membrane inserts; 8 μm pore size). The chemoattractant factor CXCL12 was added in two different concentrations (low dose = 20 ng/ml, high dose = 80 ng/ml) to 500 μl RPMI in the lower compartment. Agents that inhibit cell migration were added directly to the cell suspension in the upper compartment (CXCR4-RB (12 μg/ml), SDC4-RB (1:500), ITGB1-RB (8 μg/ml), ACKR-RB (10 μg/ml)) (Supplementary Table 2). The cells were incubated for 24 hours in a standard cell culture incubator. For quantification of migratory cells, medium was carefully aspirated from the upper compartment and all non-migratory cells inside the insert were removed with cotton-tipped swabs. Next, inserts were stained in 400 μl Cell Stain Solution for 10 minutes at RT, followed by 5x washing with PBS and air-drying. Migratory cells were imaged with a light microscope under 100x magnification objective, with at least three individual fields per insert.

### Label-free secretome analysis

Protease inhibitors (Complete mini EDTA Free, Thermo Fisher, USA) were added to collected medium and supernatant concentration was achieved by ultrafiltration using MWCO protein concentrators (Thermo Fisher, USA). Proteins (50 μg) were diluted in 1% SDS, 50 mM dithiothreitol, 100 mM Tris buffer (pH 8.0), denatured for 10 min at 95°C and digested by filter-aided (Merck Millipore, USA) sample preparation for proteome analysis (FASP)^48^. Next, overnight digestion was performed at 37°C with 60 μl digest buffer containing 500 ng trypsin (Sigma Aldrich, USA) in 50 mM triethylammonium bicarbonate buffer (pH 8.5). Peptides were recovered by adding 140 μl HPLC-grade water and centrifugation of filters for 25 minutes at 14,000 g (final volume: 200 μl tryptic digest). An aliquot corresponding to 10% of the digest volume was purified by strong cation exchange StageTips^49^. The eluate (10 μl containing 500 mM ammonium acetate in 20% acetonitrile) was diluted 10x in 0.2% formic acid and injected for nanoscale LC-MS/MS analysis (4 μl). Tryptic peptides were analysed by nanoscale LC-MS/MS by a Top-12 data-dependent analysis method run on a Q-Exactive “classic” instrument (Thermo Fisher, USA)^50^ with a gradient length of 85 min. Raw data were loaded in MaxQuant (v 1.6.2.6a) for database search and label-free quantification by the MaxLFQ algorithm (Cox Mann, MCP 2014). For data processing, default parameters in MaxQuant were adopted, except for the following: LFQ min. ratio count: 2; fast LFQ: ON; quantification on unique peptides; match between runs (MBR): activated between technical replicates, not between different samples (achieved by assigning a separate sample group to each biological sample and allowing MBR each group only). For protein identification, MS/MS data were queried using the Andromeda search engine implemented in MaxQuant against the Homo Sapiens Reference Proteome (accessed in December 2019, 74,788 sequences) and the Macaca Fascicularis Reference Proteome (accessed in January 2020, 46,259 sequences).

### RNA isolation, reverse transcription PCR (RT-PCR), and quantitative real-time PCR (qRT– PCR)

Total RNA of CFs/HVPs co-culture and conditioned CFs exposed to co-culture medium was extracted using the Absolutely RNA Miniprep Kit (Agilent Technologies, USA) according to the manufacturer’s instructions and 1 μg was reverse transcribed using the High-Capacity cDNA Reverse Transcription kit (Thermo Fisher, USA). qRT-PCR was performed using 25 ng cDNA per reaction and the Power SYBR Green PCR Master Mix (Thermo Fisher, USA). Gene expression levels were assessed in three independent biological samples and normalized to glyceraldehyde-3-phosphate dehydrogenase (*GAPDH*) expression by using the comparative cycles of threshold (Ct) method (ΔCt). qRT-PCR assays were run on a 7500 Fast Real Time PCR system (Thermo Fisher, USA) and the data were processed using 7500 software v2.3. Following primers were used: GAPDH_Fw: TCCTCTGACTTCAACAGCGA; GAPDH_Rv: GGGTCTTACTCCTTGGAGGC; ROBO1_Fw: GGGGGAGAGAGAGTGGAGAC; ROBO1_Rv: AGGCTCTCCTACTGCAACCA; SLIT2_Fw: TAGTGCTGGCGATCCTGAA; SLIT2_Rv: GCTCCTCTTTCAATGGTGCT.

### Pig experiments and treatments

Pigs were sedated by intramuscular injection of ketamine (Ursotamin^®^, Serumwerk Bernburg, Germany), azaperone (Stresnil^®^, Elanco Animal Health, Bad Homburg, Germany) and atropinsulfate (B. Braun, Melsungen, Germany) and kept in full anaesthesia with mechanical ventilation by continuous intravenous application of propofol (Narcofol^®^, CP Pharma, Burgdorf, Germany) and fentanyl (Fentadon^®^, Eurovet Animal Health BV, Bladel, Netherlands). Following left lateral thoracotomy, the pericardium was opened and the anterior wall of the left ventricle was exposed. After induction of RFA injuries and injection of HVPs, the thorax was closed using multiple layers of sutures. To continuously monitor heart rate and function during the follow-up period, a cardiac event recorder (BioMonitor 2, Biotronik, Berlin, Germany) was implanted subcutaneously on the left thorax wall of the D14 group pigs. All pigs had a central venous catheter (Careflow™, Merit Medical, Galway, Ireland) inserted via the lateral ear vein that remained in place over the whole course of the follow up period.

For additional immunosuppression, 5mg/kg methylprednisolone was applied intravenously on day 1 and 2.5mg/kg on day 2 (Urbason^®^, Sanofi-Aventis, Frankfurt, Germany). Further, all pigs received a once daily oral dose of 0.2mg/kg tacrolimus (Prograf^®^, Astellas Pharma, Munich, Germany) and 20mg/kg oral mycophenolat-mofetil (CellCept^®^, Roche, Penzberg, Germany) twice daily over the whole course of the experiment. For euthanasia, pigs were placed in full anaesthesia as described above and circulation was terminated by systemic injection of potassium chloride (B. Braun, Melsungen, Germany).

All pig experiments were performed with permission of the local regulatory authority, Regierung von Oberbayern (ROB), Sachgebiet 54, 80534 München (approval number: AZ 02-18-134). Applications were reviewed by the ethics committee according to §15 TSchG German Animal Welfare Law.

### Transplantation of HVPs into porcine hearts *in vivo*

HVPs sorted by MACS with <5% TRA-1-60^+^ cells were used for transplantation experiments in pigs. Briefly, 6×10^7^ sorted HVPs were suspended in 10 μl CCM supplemented with 5 μM ROCK inhibitor on the day of transplantation. Cells were spun down after thawing, and only the cell pellet was transplanted using insulin syringe. After exposing the pig heart by lateral thoracotomy and generation of non-transmural RFA injury as described above, the cell suspension was injected approximately 1 cm apart from the injury. At injection sides, U-stitches with pericardial patches were placed and closed using a tourniquet to reduce cell loss during cell application (Extended Data Movie 2).

### Sample processing and immunofluorescence analysis

Cells on chamber slides were fixed with 4% PFA for 10 min at RT. Co-culture 3D constructs were fixed in 4% PFA for 30 min at 4°C, cryopreserved with ice-cold methanol/acetone for 10 min at -20°C and sectioned at 12 μm in Tissue-Tek O.C.T. compound (Sakura Finetek, JP). Freshly explanted pig hearts on D3, D5 and D14 after RFA injury and HVP injection were examined for macroscopic signs of tumour formation before RFA-injured areas with corresponding HVP/control injection sites were manually excised and fixed with 4% PFA for 24 hours at 4°C, followed by cryopreservation with ice-cold methanol/acetone for 30 min at -20°C and sectioning at 12 μm in O.C.T. compound.

For immunofluorescence staining, samples were washed three times with PBS, permeabilised and blocked with 0.1% Triton X-100 and 10% fetal bovine serum (FBS) for 2 hours at RT (co-culture and monolayer samples) or overnight at 4°C (*in vivo* samples).

Primary antibodies against desired epitopes (Supplementary Table 2) were incubated overnight at 4°C at the indicated dilutions in 1% FBS, 0.1% Triton X-100 in PBS. After washing five times with 0.1% Triton X-100 in PBS for 5 minutes, appropriate secondary antibodies (1:500) were added in 1% FBS, 0.1% Triton X-100 in PBS for 2 hours at RT protected from light. After washing three times for 5 minutes with 0.1% Triton X-100 in PBS, Hoechst 33258 was added at a final concentration of 5 μg/mL in PBS for 15 minutes at RT. After washing three times with PBS, samples were covered with fluorescence mounting medium (Dako, USA) and stored at 4°C. Images were acquired using a DMI6000-AF6000 or SP8 confocal laser-scanning Leica microscope. Images were assigned with pseudo-colors and processed with ImageJ. Quantification of scar volumes, eGFP^+^ area and cell-type proportions were performed with image J cell counter notice and volume calculator.

### Statistical analysis

Statistical analyses – excluding scRNAseq experiments – are presented as mean ± SEM unless otherwise indicated. Two groups were compared using an unpaired *t*-test. Statistical tests were performed on the online platform socscstatistics.com. Significance was defined as *p ≤ 0.05, **p ≤ 0.01, ***p ≤ 0.001.

### Data and CODE availability

The RNAseq data generated during this study are available at the GEO database with project number GEO: GSE153282. The mass spectrometry data have been deposited to the ProteomeXchange Consortium via the PRIDE database with the dataset identifier PXD019521.

## Acknowledgements

We thank Drs. Rabea Hinkel (Deutsches Primatenzentrum Göttingen), Mats Spångberg, Bengt Eriksson, Astrid Fagreaus, and Pia Ekeland (Karoliska Institute), as well as Jan-Michael Abicht and Matthias Längin (Walter Brendel Centre of Experimental Medicine, LMU) for providing NHP hearts, Dr. David Elliott for his generosity of the two NKX2.5-eGFP cell lines, Dr. Alexander Goedel, Dr. Ran Yang, and Dr. Xidan Li for FACS and bioinformatics assistance, and Dr. Steffen Dietzel (Core Facility Bioimaging, Biomedical Center, LMU) for 2-Photon live microscopy. We would like to acknowledge Birgit Campbell and Christina Scherb for their technical assistance in cell culture and Sarah Luger (moments-of-aha.com) for graphic illustrations. This work was supported by grants from: the European Research Council, ERC 743225 (to K.R.C.) and ERC 788381 (to A.M.); the German Research Foundation, Transregio Research Unit 152 (to A.M., K-L.L.) and 267 (to A.M., K-L.L., and C.K.); Swedish Research Council Distinguish Professor Grant (to K.R.C.); DZHK (German Centre for Cardiovascular Research).

## Author contributions

K-L.L, A.Mo., and K.R.C. together setup the collaboration and conceived the overall experimental plan. C.S. and M.T.D.A. performed functional experiments and histochemistry, analysed data, and generated figures. K.S.F. produced HVPs and CMs, performed MACS sorting, and analysed data. C.S., M.T.D.A., and K.S.F. contributed to the conception and design of experiments. G.S. performed bioinformatics analyses. F.R. and P.H. established RFA and measured scar parameters. T.D. conducted FACS analysis. T.D., I.M., and A.Me. produced heart slices and performed cellular seeding. Y.L.T. contributed to bioinformatics. S.S. developed cellular bioprinting. K.L., R.T., and A.D. introduced and adapted biomimetic slice culture. D.S. conducted calcium imaging and analysis and analysed time-lapse. E.P., M.G., and G.C. executed mass spectrometry. A.B., N.H., M.K., and C.K. performed *in vivo* pig experiments. M.K. injected HVPs *in vivo*. V.J. performed CD68 immunodetection and analysis. N.K. generated and provided transgenic LEA29Y pigs. N.K., C.K. and A.B. supervised *in vivo* studies and provided conceptual advice. A.Mo., K.R.C., and K.-L.L. conceived and supervised this study and provided financial support. A.Mo., K.R.C., and K.-L.L. wrote the manuscript. All authors commented on and edited the manuscript.

## Competing interests statement

The authors declare that they have no competing financial interests. K.S.F. and K.R.C. are co-inventors on a patent based on the HVP technology and its applications. The HVP intellectual property is assigned to SWIBCO, a Swedish holding company.

**Extended Data Figure 1.**
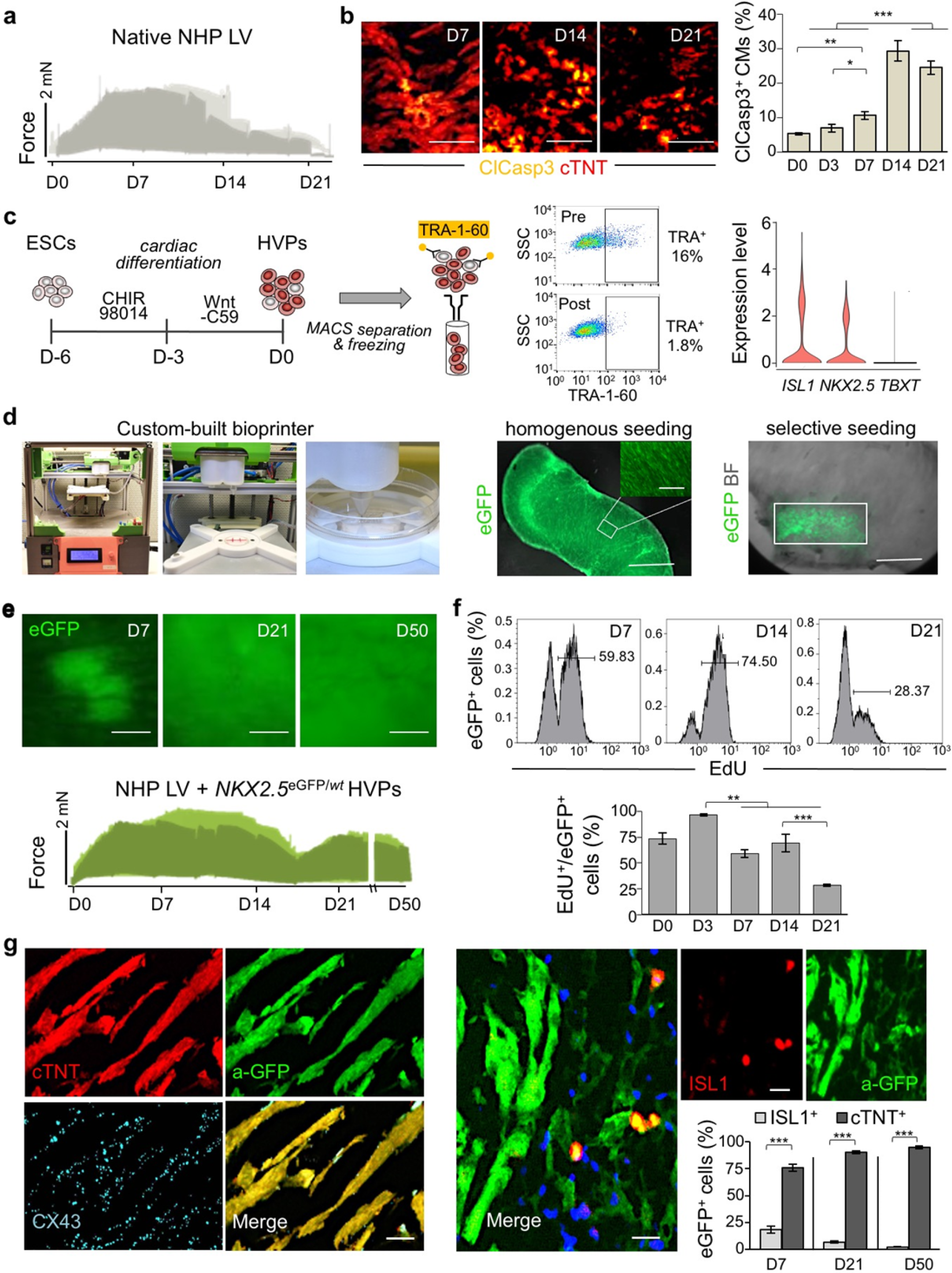
Generation and analysis of an *ex vivo* 3D chimeric human-NHP heart model. **a**, Representative, overlapped traces of contractile force of native NHP heart slices cultured *ex vivo* for 21 days in biomimetic chambers. **b**, Representative immunofluorescence images for activated cleaved caspase 3 (ClCasp3) and cardiac troponin T (cTNT) in *ex vivo* cultured NHP heart slices (left) and correspondent quantification (right) at the indicated days. Scale bars 50 μm. n ≥ 3 samples/time point. **c**, Schematic of ESC differentiation into HVPs by Wnt pathway modulation followed by MACS depletion of Tra-1-60^+^ cells and cryopreservation until seeding. Single cell RNAseq confirmed expression of *ISL1* and *NKX2.5* and loss of brachyury T (*TBXT*) on D0. **d**, Left, custom-built bioprinting device with pneumatic printhead. Right, exemplary images of homogeneous or selective seeding of eGFP^+^ HVPs onto NHP heart slices by bioprinting. Scale bars 250 μm, inlet 75 μm. **e**, Live eGFP imaging of NHP heart slices after *NKX2-5*^eGFP/*wt*^ HVP seeding at the indicated days of co-culture (top) and representative contractile force traces (bottom). **f**, Flow cytometry analysis for EdU in eGFP^+^ cells isolated at the indicated days of co-culture. n=3 samples/time point. **g**, Immunostaining of eGFP in combination with cTNT and Connexin-43 (CX43) (left) or ISL1 (right) on D50 of co-culture. Scale bars 25 μm. Bar graph shows the percentage of eGFP^+^ cells expressing ISL1 and cTNT at the indicated days of co-culture. All statistical data are shown as mean ± SEM; *p < 0.05, **p < 0.005, ***p < 0.001 (*t*-test).

**Extended Data Figure 2.**
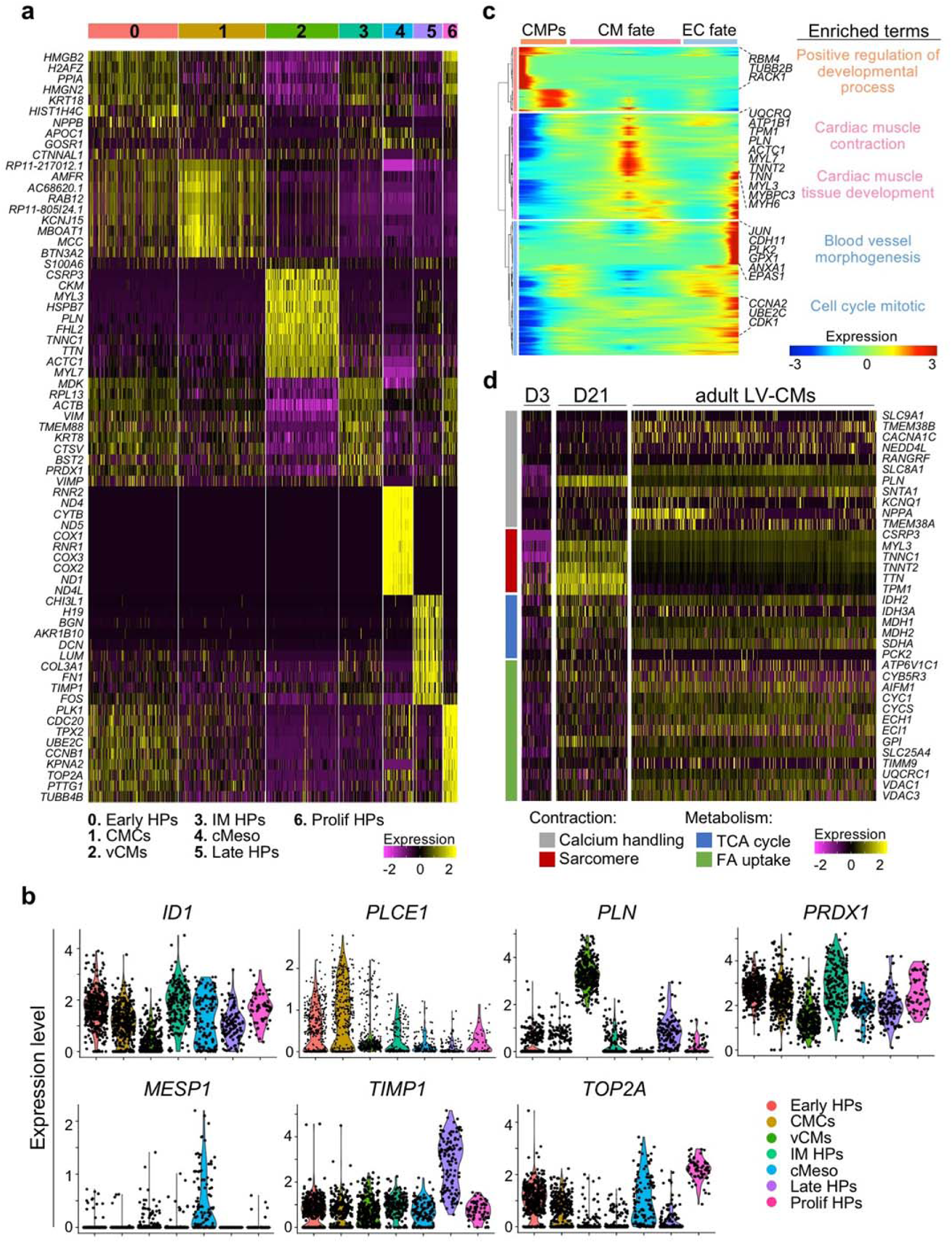
scRNAseq analysis of human *NKX2.5*^eGFP/*wt*^ HVPs in a chronic injury model of NHP heart slices. **a**, Heatmap showing expression of the top 10 genes in each cluster defined as 0- early heart progenitors (Early HPs), 1- cardiac mesenchymal cells (CMCs), 2- ventricular cardiomyocytes (vCMs), 3- intermediate heart progenitors (IM HPs), 4- cardiac mesodermal cells (cMeso), 5- late heart progenitors (Late HPs), 6- proliferating heart progenitors (Prolif HPs). **b**, Violin plots of cluster specific marker genes, *p-*value*<*0.05. **c**, Heatmap of different blocks of DEGs along the pseudotime trajectory and representative genes in each cluster. Cardiac mesodermal precursors (CMPs, D-3), endothelial cell (EC) fate (D0 and D3) and CM fate (D21). Selected top biological process and canonical pathway terms related to corresponding DEGs. **d**, Heatmap showing the expression of genes related to contraction (gray and red) and metabolism (blue and orange) in eGFP^+^ cells on D3 and D21 of *ex vivo* co-culture compared to adult human LV-CMs (Wang *et al.*, 2020). Expression levels are presented as a colour code.

**Extended Data Figure 3.**
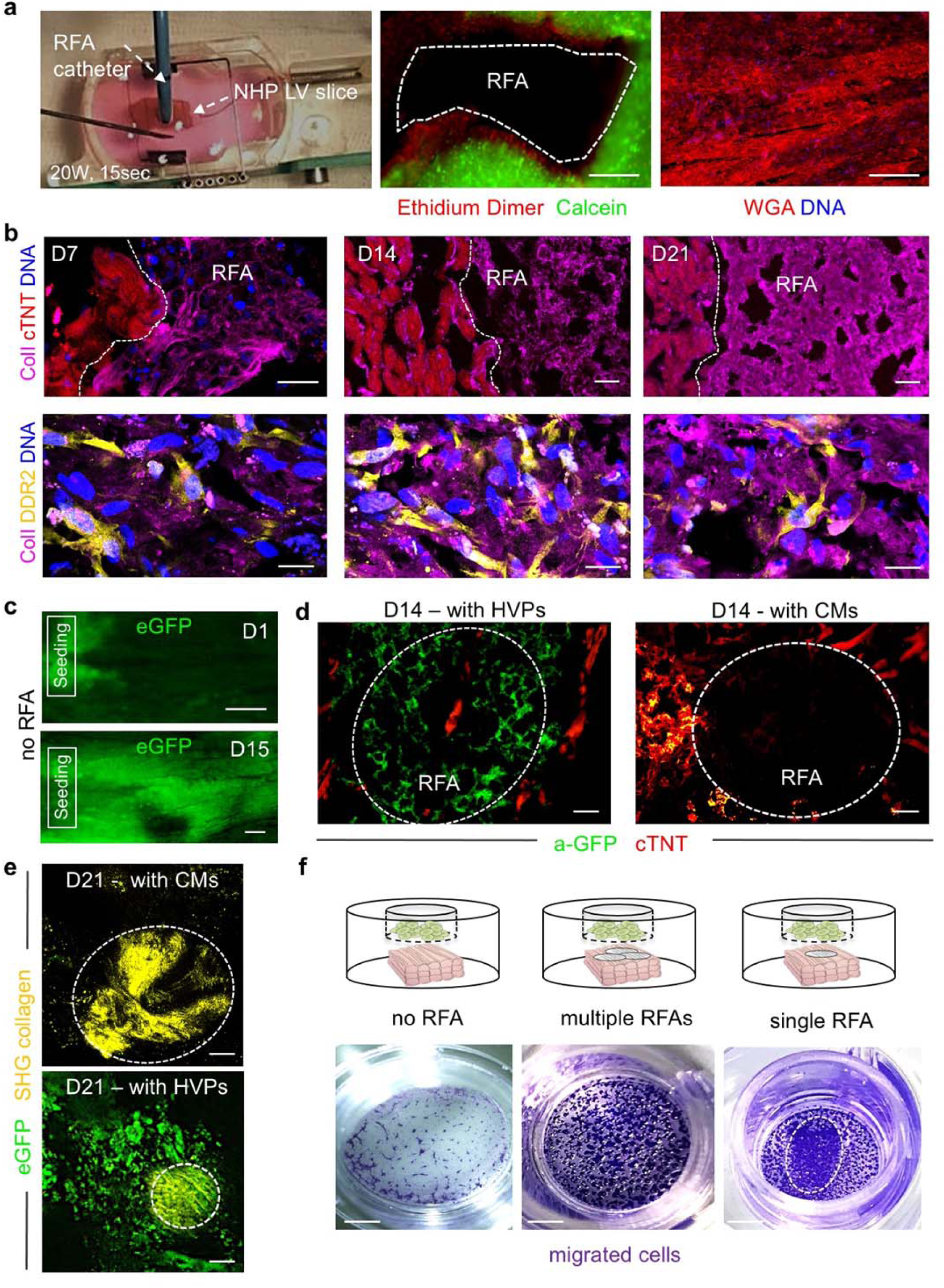
Generation and analysis of an acute *ex vivo* NHP heart injury model. **a**, Standardized non-transmural myocardial injury in NHP heart slices by defined RFA. Live and dead cells are stained by calcein and ethidium dimer, respectively (middle). ECM fibers are labeled by WGA (right). Stainings were performed immediately after RFA. Scale bar 200 μm. **b**, Representative fluorescence images of RFA-injured slices after immunostaining for Collagen type I (ColI) combined with cTNT (top) or DDR2 (bottom) on indicated days. Lower panels show images of the RFA area. Scale bars 30 μm (top) and 25 μm (bottom). **c**, Sequential live imaging of *NKX2-5*^eGFP/*wt*^ HVPs migrating from the seeding frame into the tissue showing homogenous repopulation of the slice by D15 in the absence of RFA injury. Scale bars 200 μm. **d**, Representative immunostaining of eGFP and cTNT in RFA-injured area on D14 after selective seeding of *NKX2-5*^eGFP/*wt*^ HVPs (left) or CMs (right). Scale bars 50 μm. **e**, Two-photon live microscopy of RFA-injured slices for eGFP and second-harmonic-imaging (SHG) visualization of collagen and scar size on D21. Circles demarcate areas with collagen deposition. Scale bars 100 μm. **f**, Trans-well migration assays with D0 *NKX2-5*^eGFP/*wt*^ HVPs in the upper and NHP heart slice in the lower compartment, respectively. Images show trans-well migrated HVPs on polycarbonate membrane in the absence (left) or presence of multiple (middle) or single (right) RFA injury. Dashed line marks the site of HVP accumulation. Scale bars 2 mm.

**Extended Data Figure 4.**
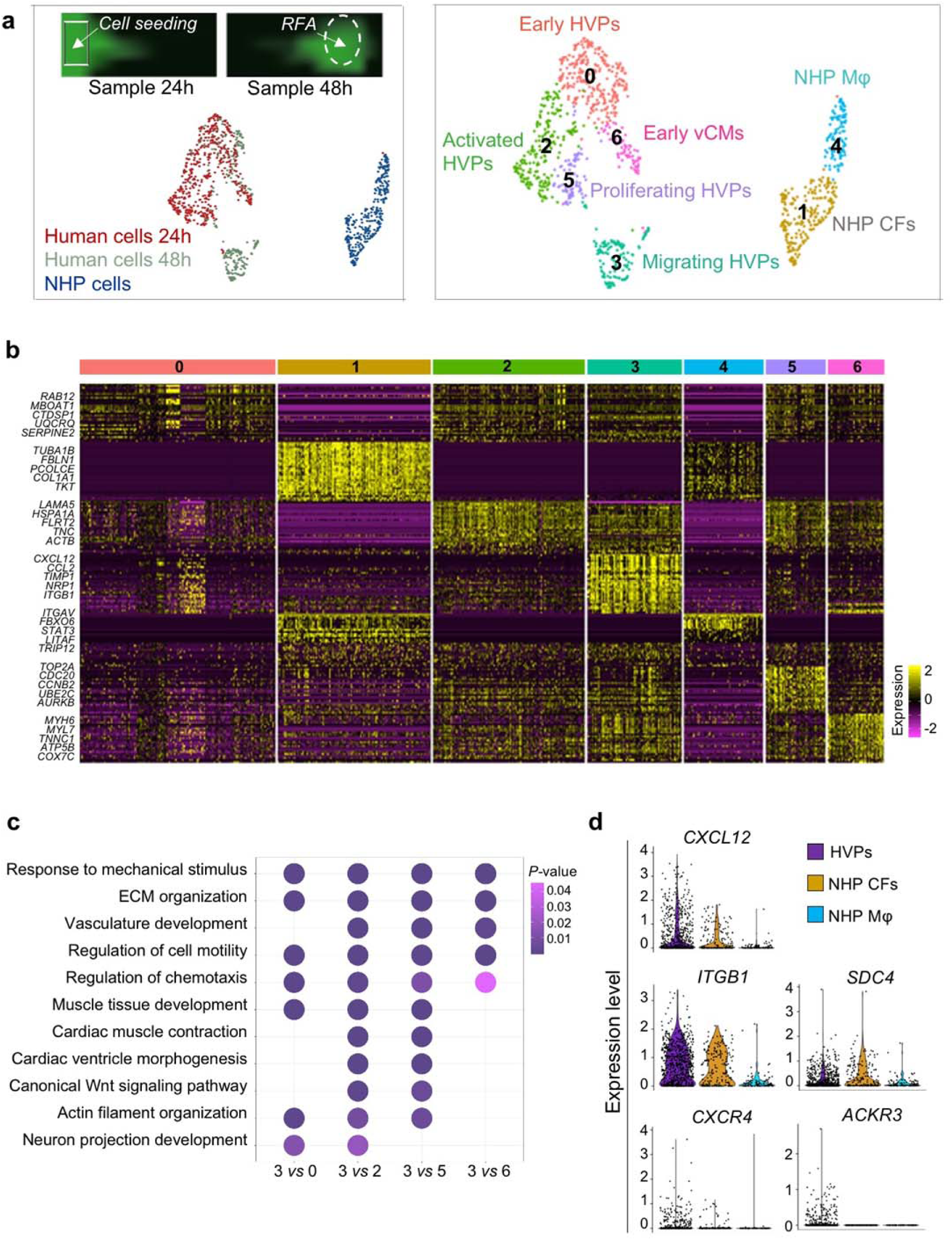
scRNAseq analysis of human *NKX2.5*^eGFP/wt^ HVPs and NHP cardiac cells after acute RFA heart injury. **a**, Left, representative images of HVPs seeded on an injured NHP heart slice at the time points used for cell collection (24h and 48h) (top) and UMAP plot of all captured cells (bottom). Right, relative UMAP clustering of captured cells. NHP, non-human primate; RFA, radiofrequency ablation; HVPs, human ventricular progenitors; vCMs, ventricular cardiomyocytes; Mφ, macrophages. **b**, Heatmap of top 50 genes in each cluster with representative genes indicated. **c**, Representative GO terms upregulated in cluster 3 (migrating HVPs) compared to the other human clusters (0, 2, 5, 6). **d**, Violin plots of *CXCL12* and its binding targets in HVPs, NHP CFs and NHP Mφ.

**Extended Data Figure 5.**
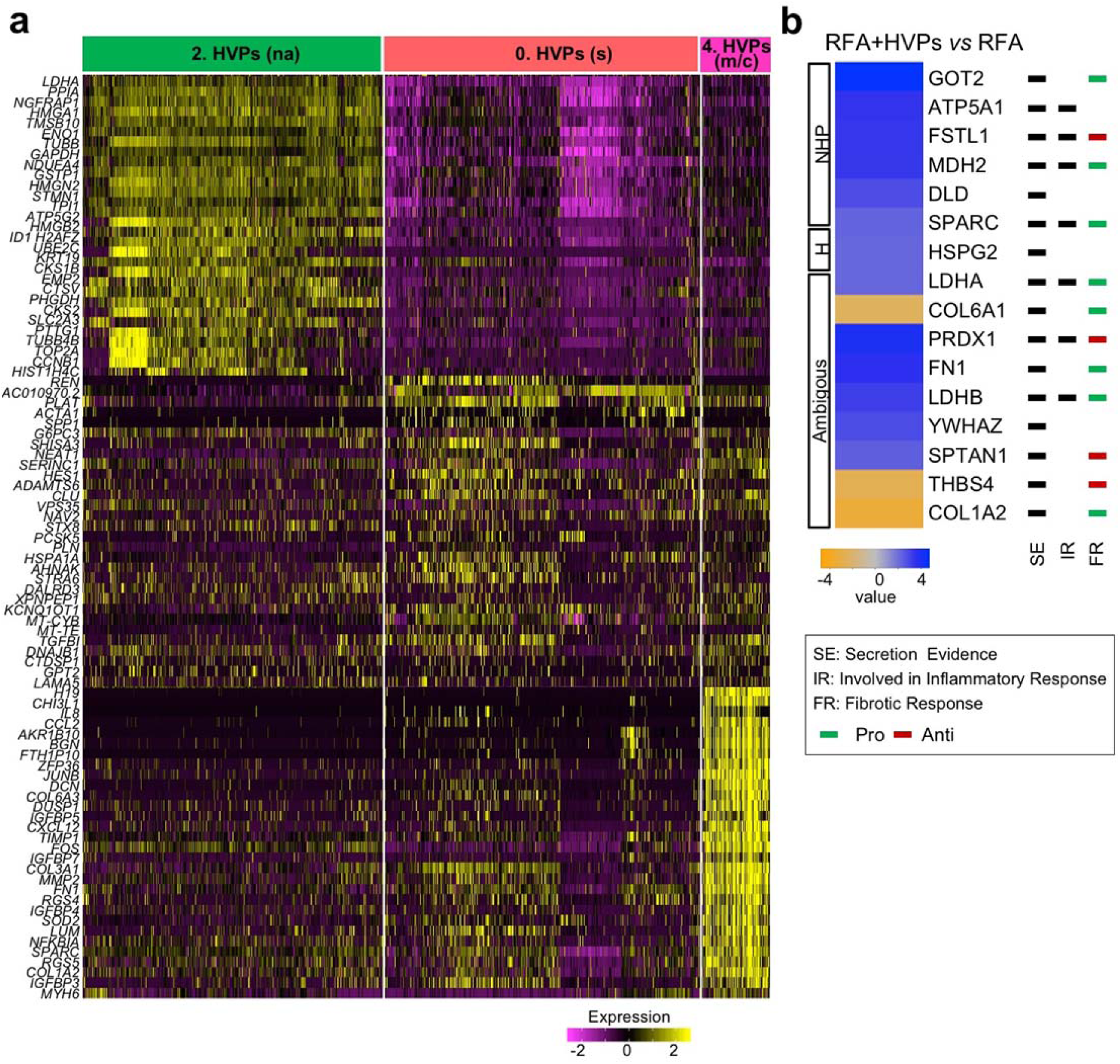
Gene signatures of dynamical cardiac progenitor states and proteomic analysis of secretome during acute injury response. **a**, Heatmap of top 30 genes depicting the expression of DEGs in non-activated HVPs (cluster 2), sensing HVPs (cluster 0), and migrating/counteracting HVPs (cluster 4). **b**, Proteomic analysis of supernatant of injured NHP heart slices with and without application of HVPs at 48h after RFA. NHP, H, and ambiguous, was assigned to proteins for which the majority of identified peptides belonged to protein sequences of macaca fascicularis, homo sapiens, or both species, respectively. n=3 biological replicates per group, *p-*value ≤0.05.

**Extended Data Figure 6.**
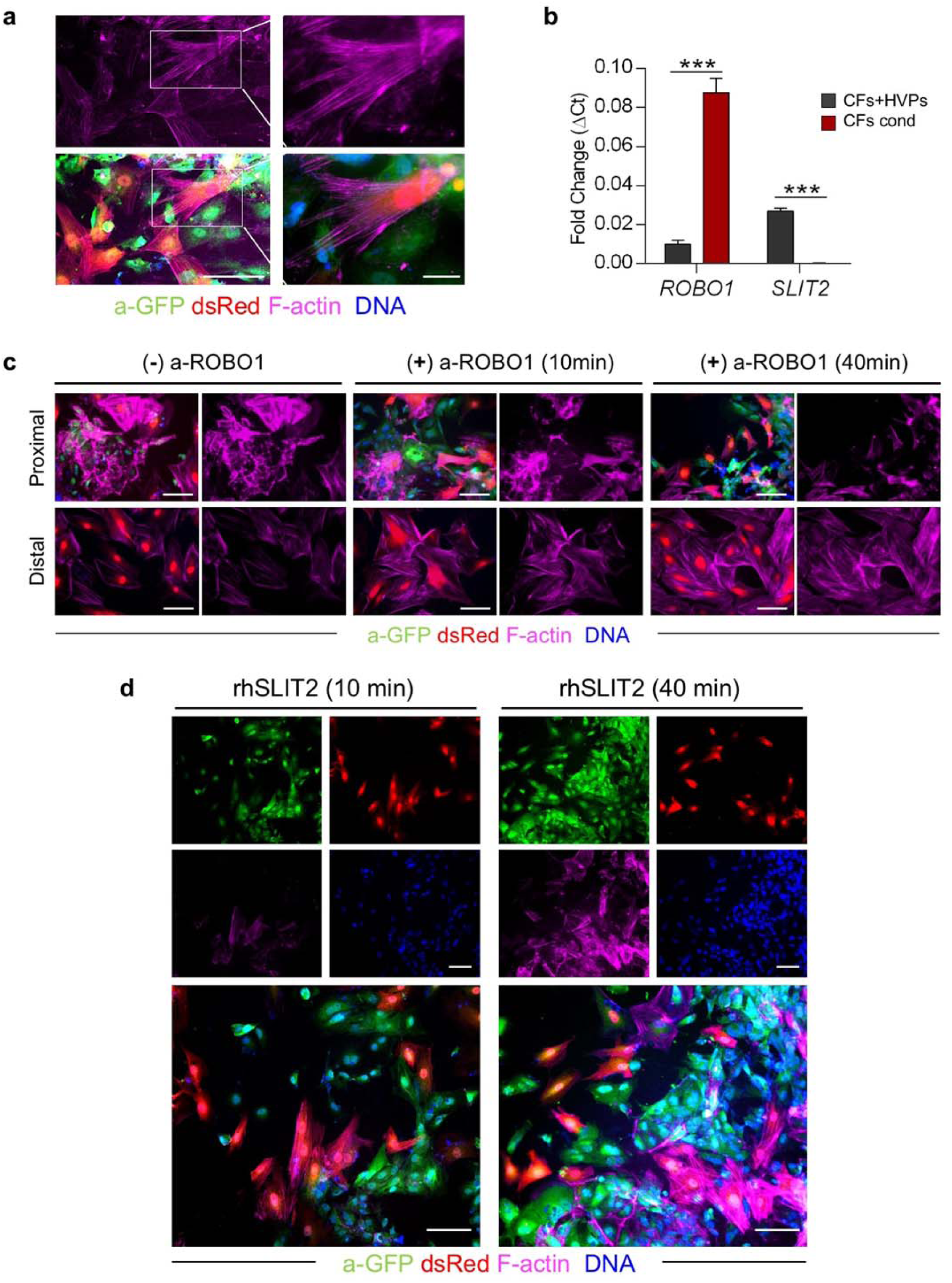
Analysis of CF repulsion signaling during acute injury response in 2D monolayer. **a**, Representative images of eGFP^+^ HVPs and dsRed^+^ CFs after F-actin staining during the repulsion phase in the injury area on D8. **b**, Quantitative RT-PCR analysis of *ROBO1* and *SLIT2* expression in injured CFs cultured with HVPs (CFs+HVPs) or alone in conditioned medium from HVP-CF co-culture (CFs cond) on D8. Data are mean ± SEM, n=2. \*\*\**p*<0.001 (*t*-test). **c**, Representative F-actin immunostaining on D8 in standard condition and after ROBO1 antibody exposure for 10 and 40 minutes showing CFs in contact with HVPs (proximal) and CFs in the remote area from the injury site (distal). **d**, Immunodetection of eGFP in conjunction with Phallodin (F-actin) stain in HVP-CF co-culture on D8 after recombinant human SLIT2 (rhSLIT2) exposure for 10 and 40 minutes. Nuclei were counterstained with Hoechst and CFs are labelled with dsRed (**a, c, d**). Scale bar, 75 μm (**a, c, d**).

**Extended Data Figure 7.**
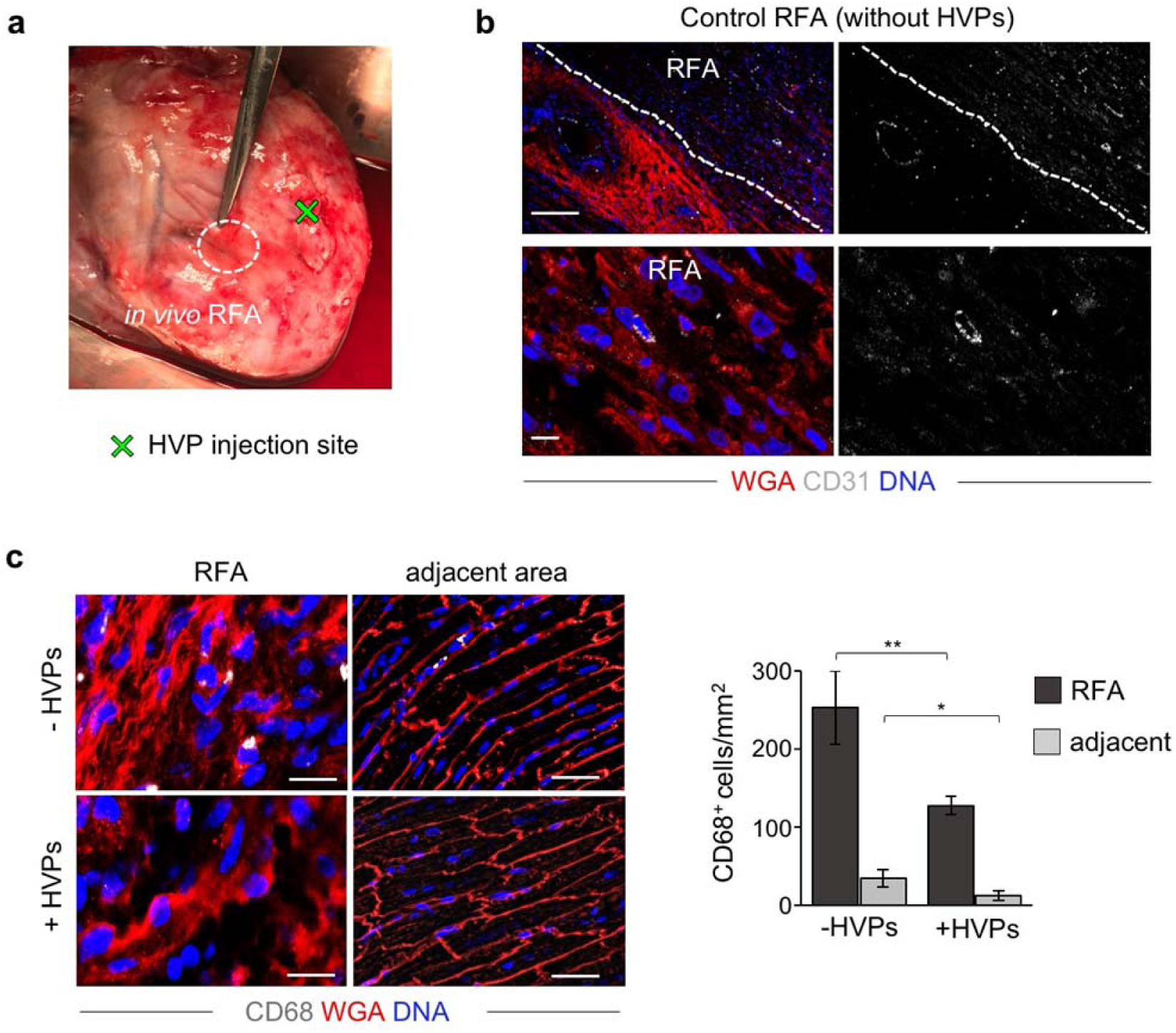
Macro- and microscopic analyses of LEA29Y pig hearts after RFA injury and HVP injection *in vivo.* **a**, Image of a freshly explanted LEA29Y pig heart 14 days after *in vivo* RFA and adjacent HVP injection showing no macroscopic signs of teratoma formation. **b**, Representative fluorescence images of control RFA and adjacent area (top) or magnified zoom of control RFA (bottom) after CD31 immunodetection and WGA labelling. Scale bars 100 μm (top) and 10 μm (bottom). **c**, Representative immunofluorescence stainings of CD68 (left) and correspondent statistical analysis (right) in RFA and adjacent areas in the presence and absence of HVPs. Scale bars 25 μm. Data are shown as mean ± SEM, n=7 slices from 2 pigs per group. *p<0.05, **p<0.005 (*t*-test).

## Supplementary Information

**Supplementary Table 1.** Data source for scRNAseq and proteomic analyses

**Supplementary Table 2.** List of antibodies, fluorescent probes, recombinant proteins, and assays used in the study

**Extended Data Movie 1.** Time-lapse live imaging of HVPs (green) and NHP CFs (red) at the injury site (frame time: 90 minutes, duration: 3 days)

**Extended Data Movie 2.** RFA and cell transplantation procedure in LEA29Y porcine hearts

